# Effects of spectral light quality on the growth, productivity, and elemental ratios in differently pigmented marine phytoplankton species

**DOI:** 10.1101/2020.04.09.034504

**Authors:** T. L. Bercel, S. A. Kranz

## Abstract

Effects of light quality on the growth, productivity, and cellular composition of three uniquely pigmented marine phytoplankton species were characterized. To accomplish this, cultures of *Prochlorococcus marinus, Synechococcus sp*., and *Thalassiosira weissflogii* were grown under three commercially available LEDs as well as a fluorescent growth light. Despite having equal photosynthetically active radiation, light quality and thus photosynthetically usable radiation differed between the treatments. Growth was unaffected in all species tested, yet primary productivity was affected in *P. marinus* and *Synechococcus sp.* All species regulated cellular carbon and nitrogen quotas as a direct response to light spectra, while cellular chlorophyll *a* was regulated in *Synechococcus sp.* and *T. weissflogii* only. Analysis of pigment ratios revealed minor acclimations in some of the cultures and photophysiological analysis indicated changes in the photoacclimation state between different light environments. These results show that while the species used in our experiment are able to maintain growth when exposed to lights of varying quality, underlying cellular metabolism and biochemistry can be affected. The data presented here highlight the importance of carefully choosing a lighting environment with a defined spectral quality when designing laboratory-based experiments or setting up bioreactors for biomass generation.

**Highlight:** With light emitting diode-based growth lights becoming available to researchers, it is important to consider the spectral quality of light when designing experiments to understand responses of phytoplankton to environmental conditions.

## Introduction

Marine phytoplankton are mainly photoautotrophic organisms. As such, phytoplankton require the ability to efficiently convert light energy into biochemical energy via the process of photosynthesis. Thus, light availability is a key environmental factor affecting phytoplankton growth, productivity, and niche distribution (Boyd et al., 2010; Six et al., 2007). Numerous studies have been conducted investigating the effect of light intensity and/or duration on phytoplankton physiology (e.g. Brand and Guillard, 1981; Falkowski, 1984; Falkowski and Owens, 1978; Glover et al., 1987; Hessen et al., 2008; see Lehmuskero et al., 2018 for review). Furthermore, when other environmental parameters, such as temperature, CO_2_, salinity, and/or nutrient concentrations are varied, the availability of light (both the intensity of the irradiation and the duration) affect the ecophysiological responses of the phytoplankton cultures or communities (Boatman et al., 2017; Dickman et al., 2006; Feng et al., 2008; Finkel et al., 2006; Gao et al., 2012; Healey, 1985; Ivanikova et al., 2007; Kranz et al., 2010; Redalje and Laws, 1983; Rhee and Gotham, 1981; Rost et al., 2006). In addition, it has been shown that a dynamically changing light intensity or simply changes in daylength can affect the cellular response and energetics of phytoplankton (Hoppe et al., 2015; Rost et al., 2006; White et al., 2020).

Therefore, when aiming to better understand and predict phytoplankton responses to environmental changes using quantitative data on cellular response parameters such as growth, productivity, and elemental composition, it is imperative to define the light conditions well. To identify or even separate the complex correlations between cellular responses and environmental parameters, scientists often use controlled laboratory environments, following best practice guides (e.g. Riebesell et al., 2011 and others). We, however, often ignore that the light quality (light spectrum) in the laboratory does not always mimic that of the natural environment in which phytoplankton thrive.

Historically, laboratory studies investigating the physiological responses of marine phytoplankton species to environmental change have utilized ‘daylight’ or ‘growth light’ fluorescent bulbs to supply light for growth. These specific fluorescent bulbs have an emission spectrum that usually resembles that of the natural solar light experienced at the surface of the ocean (e.g. GE Daylight Ecolux^®^ T12 fluorescent light bulbs). However, a disadvantage of using fluorescent bulbs is that they cannot easily replicate the dynamic and occasionally extreme high light intensity experienced by phytoplankton in nature, nor can their emission spectra be easily modified. Recently, light emitting diode (LED) based light incubators and stand-alone growth lights have become readily available. As LED systems have become more flexible and powerful, LED light incubators have also become more commonly utilized in culture studies of phytoplankton (see Schulze et al., 2014 for a review). One of the disadvantages of LEDs is, however, that their spectra are typically restricted to narrow emission peaks and hence a mixture of two or more LEDs are used to cover the major light spectra needed for growth of photoautotrophic cells. In order to understand wavelength specific responses of phytoplankton, multiple studies have been performed using different LED systems to quantify specific photosynthetic responses to distinct wavelengths (Abiusi et al., 2014; Kim et al., 2013; Marchetti et al., 2013; Miao et al., 2012; Rendon et al., 2013; Schellenberger Costa et al., 2013; Stadnichuk et al., 2011; Wang et al., 2007; Xu et al., 2013). However, the effects of LED systems designed to mimic natural spectra are sparsely investigated (Göritz et al., 2017). While the laboratory settings provide a controlled environment for experiments, they usually have a limited capacity to simulate ecologically relevant light regimes which in turn limits the accuracy of projecting the findings to the real world.

To better understand why growth light spectra matters it is important to recognize that all phytoplankton species contain a specific set of pigments. In general, phytoplankton pigments can be thought of as photoactive or photoprotective. Photoactive pigments’ absorption contributes to energy capture which leads to electron transport and ATP and NADPH production, which are then used as energy for carbon fixation and other metabolic processes. Photoactive pigments include chlorophylls (like chl *a)* as well as accessory pigments, like xanthophylls or those found in cyanobacterial phycobilisomes. Photoprotective pigments can absorb excess light energy previously absorbed by chl *a* and dissipate it as heat or fluorescence, thus avoiding photodamage to the reaction centers in the photosystem. Carotenoids are generally thought of as the main photoprotective pigments found in algae and higher plants, however some such as fucoxanthin can be photosynthetically active (Wilson et al., 2008). Since the composition of photoactive and photoprotective pigments is species specific and as species can alter the relative contribution of pigments depending on light and nutrient availability, it is evident that the choice of growth light will affect phytoplankton in many, yet unknown ways. The goal of this study is to characterize physiological responses of three uniquely pigmented phytoplankton species to different light systems which could be used for phytoplankton culturing. To accomplish this, phytoplankton were cultured under traditionally used fluorescent light as well under three different LED systems with unique emission spectra. Cellular responses, including growth, productivity, elemental composition, chl *a* content, spectral pigment deconvolution, and photophysiological properties were analyzed.

## Materials and Methods

### Culture conditions

*Prochlorococcus marinus* (CCMP1986), *Synechococcus sp.* (CCMP1334), and *Thalassiosira weissflogii* (CCMP1336) obtained from the National Center for Marine Algae and Microbiota were acclimated to four different light sources using a semi continuous batch culture approach. All cultures were maintained at 22°C in a walk-in temperature controlled room and grown in modified artificial seawater based on Aquil media (Price et al., 1989) with 25 μM Na_2_HPO_4_ as phosphorus source as well as 400 μM NaNO_3_ as nitrogen source for *Synechococcus sp.* and *T. weissflogii* and 400 μM NH4Cl as nitrogen source for *P. marinus.* Additionally, *P. marinus* only received thiamine for vitamins and 1/10^th^ of the typical Aquil metal mix added. *T. weissflogii* was supplemented with 50 μM Si for growth. *T. weissflogii* and *P. marinus* were maintained at 100 *μ*mol photons m^-2^ s^-1^ while *Synechococcus sp.* was maintained at 50 *μ*mol photons m^-2^ s^-1^. Light was on a 12:12 hr light:dark cycle and illumination intensity remained constant during the photoperiod. Cultures were grown and tested in triplicate and given at least 7 generations (approximately 2-4 weeks) to adapt to the different light sources before being used for experiments.

Lights used for culture growth include a Fluorescent bulb (GE Daylight Ecolux^®^ T12), a Bright White LED (Bright White LED Strip Lights, Cool White), an Aquarium LED (NICREW ClassicLED Plus LED Aquarium Light), and a phytoplankton growth light (Kyocera Aqua light). Each light source has a unique light emission spectrum, which is shown in Figure 1 A-D.

**Figure 1:**
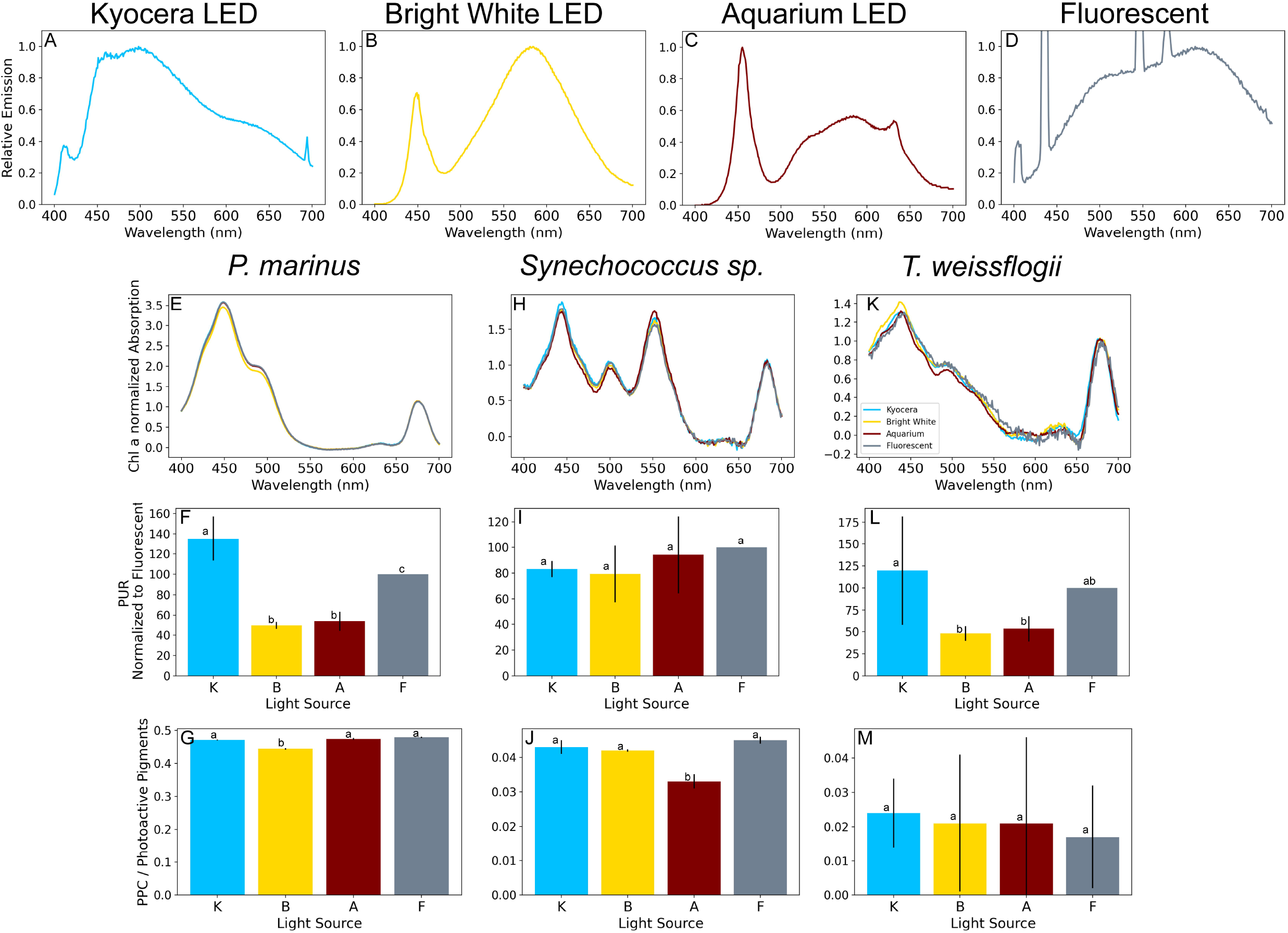
Emission spectra in photons m^-2^ for the light treatments used in this study (A-D). Peaks shown in the fluorescent spectrum (D) are cut off. Whole cell light absorption normalized to mg Chl *a* L^-1^. PUR estimates shown normalized to Fluorescent acclimations, and PPC/Photoactive ratios obtained from spectral fitting for *Prochlorococcus marinus* (E-G), *Synechococcus sp.* (H-J), and *Thalassiosira weissflogii* (K-M). Values shown are mean values for biological replicates ± s.d, n ≥ 2.

### Growth, Chl *a*, Elemental composition, Absorbance

Specific growth rates were measured via cell count using a flow-cytometer (CytoFLEX (Beckman, Indiana, USA)) and chl *a* fluorescence using a fluorometer (Trilogy, Turner Design, California, USA). The specific growth rate was estimated by an exponential fit through the data during exponential growth. All experiments and analytical samplings were performed during mid exponential growth. Samples for cellular chl *a* concentration were taken using gravity filtration onto a GF/F filter. The filter was stored at −20°C until processing. Chl *a* was extracted in 90% acetone for 24 hours in the dark at −20°C, sonicated for a brief period and subsequentially measured using a UV/VIS spectrophotometer (Evolution 220, ThermoFisher, Massachusetts, USA). Chl *a* concentration was calculated following Eq. 1, adapted from the JGOFs protocol:

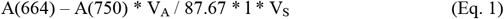

Where A(664) and A(750) are the absorbance at 664 and 750 nm, V_A_ is the volume of acetone used for extraction in L, 87.67 is the extinction coefficient for chl *a* in 90% acetone, l is the cuvette length in cm, and V_S_ is the volume of sample filtered. Particulate organic carbon and nitrogen (POC and PON) samples were taken in the same manner. For POC and PON precombusted GF/F filters (400°C, 12hr) were used, followed by an acidification and drying step before sending the compressed filters to the Stable Isotope Facility at UC Davis for analysis via a PDZ Europa ANCA-GSL elemental analyzer interfaced to a PDZ Europa 20-20 isotope ratio mass spectrometer (Sercon Ltd., Cheshire, UK).

Cellular light absorbance spectra were obtained by measuring the absorbance of a midexponential culture sample in artificial seawater media between 400-800 nm at 1 nm increments in a UV/VIS spectrophotometer using a 10 cm quartz cuvette (Evolution 220, ThermoFisher, Massachusetts, USA). In order to determine the contribution of individual pigments to the absorption spectra, a spectral deconvolution method based on Thrane et al., (2015) was applied. It is important to note that within this deconvolution method, a background spectra which accounts for light attenuation not attributable to pigment absorption (e.g. light scatter) is accounted for (see Fig S1 – S3). *In vivo* pigment absorption spectra were used as described in Bidigare et al. (1990) and adjusted for absorbance peak shifts which occur due to protein-pigment complexes which form *in-vivo* (Prézelin, 1981). Parameterization used for our study are given for each species and the respective pigments in Table S1. Additional information on the model used is given in the supplemental material.

Determination of photosynthetically usable radiation (PUR) was calculated using Eq. 2:

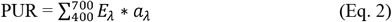

where E_λ_ is the relative quantity of photons emitted (normalized to the maximum emission peak) at each wavelength λ and a_λ_ is the chl *a* normalized % absorbance for each phytoplankton culture wavelength λ (Morel 1978). Emission spectra for the different growth lights were measured using a calibrated FLAME-S-VIS-NIR-ES Spectrometer (Ocean Insight). Light emission at each wavelength (given in Watts/nm) was further converted into photon flux at each wavelength.

### ^14^C Primary Productivity

Rates of primary productivity (PP) were determined using ^14^C labeled bicarbonate (NaH^14^CO_3_) (Nielsen, 1952). Cultures in mid-exponential phase were placed in 20 mL borosilicate glass reaction vials and spiked with 0.5 μCi of NaH^14^CO_3_. Cultures were then placed in their respective growth chambers. Measurements started at the beginning of the dark period to allow for bottle acclimation to occur in the dark. Based on this setup, estimates of productivity (further referred to as PP) have to be assumed to be between net and gross primary production. Over the duration of the light phase, vials were inverted regularly to avoid settling. A dark and t=0 control were performed for reference. Incubations were stopped after 24 hr at the beginning of the following dark period by filtration with gentle pressure onto either GF/F (*T. weissflogii)* or 0.22 μM PES filters (*P. marinus and Synecho<coccus sp*.). Filters were placed into 20 mL scintillation vials and 600 μL of 6N HCl was added to remove all remaining ^14^CO_2_. After 24 h of degassing, 10 mL of scintillation cocktail was added to each filter. Samples were counted in a Liquid Scintillation Counter (PerkinElmer TriCarb 5110 TR) to obtain disintegrations per minute (DPM). The total ^14^C spike was determined using an aliquot of each sample at the time of incubation termination, transferred into 50 μL of ethanolamine, and 7 mL of scintillation cocktail was immediately added. Rates of PP were then calculated according to Eq. 3:

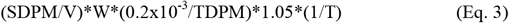

Where SDPM is the sample DPM, V is the volume of the sample in L, W is the concentration of dissolved inorganic carbon in the incubations (around 2300 μM), 0.2×10^-3^ is the volume of the total count aliquot in L, TDPM is the sample’s total DPM count and 1.05 represents the isotope discrimination factor between ^12^C/^14^C. T is time in hours. Rates of PP were subsequently normalized to the cellular chl *a* content determined prior to the incubation.

### Oxygen Evolution

Rates of photosynthetic oxygen evolution were measured using a FirestingO2 optical oxygen meter and respiration vials (Pyro Science, Germany). Cultures in mid-exponential phase placed in front of their respective light source at acclimation light intensity. Care was taken that no headspace was left, and a thin syringe needle was injected to allow for potential pressure adjustment. The cultures were continuously mixed using magnetic stir bars. Oxygen evolution was measured in the cultures for 24 hours undisturbed in front of their growth lights. An abiotic control containing fresh sterile, filtered seawater was also measured for blank correction. Net oxygen evolution rates were determined with linear regressions of the data during the light phase. O_2_ evolution was subsequently normalized to an average cellular chl *a* determined prior to and after the experiment.

### Photophysiological parameters

Rapid fluorescence induction light curves (FLC) were measured using a Fast Repetition Rate Fluorometer (FRRf, FastOcean PTX, Chelsea Technologies Group) along with a FastAct Laboratory system (Chelsea Technologies). FLCs for each species were measured within 2 hours in the middle of the photoperiod (5-7 hours after onset of the light). Cells were dark acclimated for 5 min before the start of each FLC and each of the 10 light steps lasted 1 min each. Measured fluorescence parameters (Fo, Fm, F’, Fm’), along with estimations of the functional absorption cross section area of PSII (σ), non-photochemical quenching using the normalized Stern-Volmer quenching coefficient (NSV), and primary productivity, JVPII, were obtained from the Act2Run software and their derivation can be found in Oxborough et al. (2012). To account for differences in biomass between samples at the time of FRRf measurement, JVPII data was normalized according to the following equation:

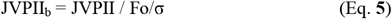

Biomass normalized JVPII_b_ was then fit using the Scipy package functions in python to obtain the descriptive Photosynthesis vs Irradiance (P vs. I) curve parameters: light saturation point (E_K_), maximum electron transport rates (Pmax), and the slope for the light limited photosynthesis (α).

## Results

### PUR, Growth, Elemental Composition, Chl *a* Content

The relative photon emission spectra for wavelengths in the photosynthetically active radiation spectrum (PAR, 400-700 nm) of the chosen growth lights are shown in Figure 1 A-D. The Kyocera LED (Fig 1A) has a main emission peak around 500 nm with significant light emission between 450 and 600 nm. Between 600 and 700 nm the emission is strongly reduced. This light was designed to mimic a natural light spectrum as seen in 5-10 m water depth of clear ocean waters. The Bright White LED (Fig 1B) has a main emission peak at 570 nm and a secondary peak at around 450 nm. A significant dip in the emission spectrum with a minimum at around 480 nm is characteristic for this kind of “off the shelf white LED” light. The Aquarium LED (Fig 1C) has a narrow peak around 450 nm and a broad peak spanning 525-650 nm, due to multiple differently colored LEDs embedded in the array. The Fluorescent bulb (Fig 1D) has a main broad peak spanning 500-650 nm, with high spikes at approximately 400, 430, 550, and 575 nm.

Phytoplankton absorption for *P. marinus* showed relatively similar patterns between the differently acclimated cultures (Fig. 1E). All but the Bright White LEDs were indistinguishable, with Bright White LED cells having a slightly reduced absorbance around 400-500 nm. Characteristic peaks for chl *a* were observed at approximately 450 and 676 nm and a peak likely representing beta-carotene in the 500 nm region. Estimates of the relative amounts of Chl *a* and *b,* as well as photoprotective carotenoids (PPC) are shown in Table S2 and indicate that 44% to 47% of the light absorption originates from non-photosynthetic active pigments. Despite the seemingly similar absorption scans, a slightly, yet, significantly lower ratio of PPC/Photoactive pigments was found in the Bright White acclimation compared to the other treatments (Figure 1G, Table S2) (One Way ANOVA, p < 0.01).

Similar to *P. marinus,* no major differences in absorption were observed in *Synechococcus sp.,* yet, in this species the Aquarium light treatment showed the lowest absorption around 400-525 nm and the highest around 525-575 nm (Fig. 1H). Characteristic peaks for chl *a*, phycoerythrin (PE, at 550 nm), and a carotenoid, most likely zeaxanthin (500 nm) were observed. Spectral deconvolution estimated the relative amounts of Chl *a*, PE, as well as PPC (Table S2), indicating that PE was around 3.5 times the amount of chlorophyll and that 3% to 4% of the total pigments are non-photosynthetically active pigments. The cultures acclimated to the Aquarium light showed a slightly, yet, statistically significant higher PE/Chl *a* ratio (Table S2) as well as a slightly lower PPC/Photoactive pigment ratio (Fig. 1J) than the rest of the cultures (One Way ANOVA, p < 0.01 for both). Despite statistical significances in the differences, these changes appear to be small.

In *T. weissflogii,* the Bright White acclimation showed a slightly higher relative chl *a* peak around 450 nm compared to the other lights (Fig. 1L). Chl *a* and *c,* as well as photosynthetically active carotenoids (PSC) and PPC were estimated in our spectral deconvolution and pigment ratios and are reported in Table S2 indicating that about 2% of the total pigments are non-photosynthetically active pigments. No significant statistical differences for pigment ratios were found between treatments (Fig 1M, Table S2) (One Way ANOVA, p > 0.05).

Based on the emission spectrum of the growth light and the absorption analysis, photosynthetically usable radiation (PUR) differed between cultures and light acclimations. In *P. marinus,* highest PUR was identified for the Kyocera acclimated cells, followed by the Fluorescent light while the Bright White and Aquarium acclimation showed strongly reduced PUR (roughly 40% of PUR calculated for Kyocera) (Fig. 1F) (One Way ANOVA, p < 0.01). In *Synechococcus sp.,* PUR was not significantly different across treatments (Fig. 1I) (One Way ANOVA, p > 0.05). In *T. weissflogii,* PUR was highest in the Kyocera and Fluorescence treatments and significantly reduced in the Bright White and Aquarium treatments (Fig. 1L) (One Way ANOVA, p < 0.01).

Growth rates were unaffected across light treatments for all species tested (Fig. 2A, E, I Table S3) (One Way ANOVA, p > 0.05). Cellular chl *a* content in *P. marinus* did not significantly change between light treatments (Fig. 2B, Table S3) (One Way ANOVA, p > 0.05) while chl *a in Synechococcus sp.* cells grown under the Fluorescent light exhibited significant higher cellular concentrations compared to the Aquarium acclimated cells (Fig. 2F, Table S3) (One Way ANOVA, p < 0.05). *T. weissflogii* also had higher chl *a* cell^-1^ in the Fluorescent acclimation compared to the Aquarium acclimation (Fig. 2J, Table S3) (One Way ANOVA, p < 0.05).

**Figure 2:**
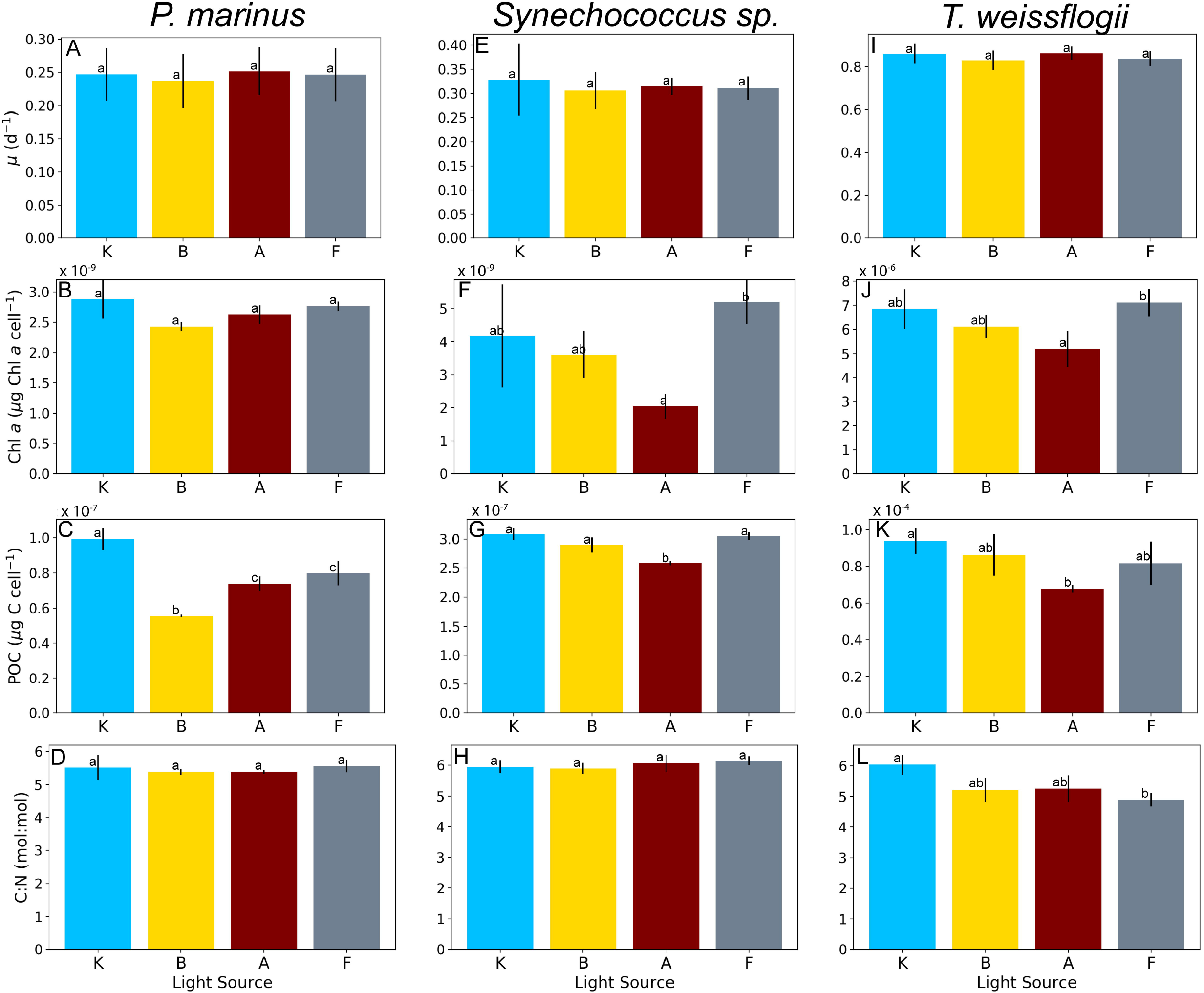
Physiological results for specific growth rate (*μ*), Chl a and POC cell^-1^ content, as well as mol:mol C:N ratios for *Prochlorococcus marinus* (A-D), *Synechococcus sp.* (E-H), and *Thalassiosira weissflogii* (I-L). Values shown are mean values for biological replicates ± s.d, n ≥ 2. Letters represent significant groupings from one-way ANOVA and HSD post hoc tests (p < 0.05).

Cellular POC and PON contents were significantly different for *P. marinus,* being highest for the Kyocera light (99.2 ± 6.2 fg C cell^-1^, 21.0 ± 1.1 fg N cell^-1^) and lowest for the Bright White (55.5 ± 0.8 fg C cell^-1^, 12.0 ± 0.4 fg N cell^-1^) with no change seen between the Aquarium and Fluorescent treatment (Fig. 2C, Table S3) (One Way ANOVA, p < 0.01, p < 0.01). C:N ratios (mol:mol) did not change significantly between treatments (Fig. 2D, Table S3) (One Way ANOVA, p > 0.05). *Synechococcus sp.* grown under the Aquarium treatment had the lowest cellular POC and PON (258.6 ± 3.7 fg C cell^-1^, and 49.8 ± 1.6 fg N cell^-1^) (Fig. 2G, Table S3) (One Way ANOVA, p < 0.01, p < 0.01). C:N ratios for *Synechococus sp.* showed no significant change (Fig. 2H, Table S3) (One Way ANOVA, p > 0.05). In *T. weissflogii* POC but not PON showed significant changes in across light treatments (Fig. 2K, Table S3) (One Way ANOVA, p – 0.05). *T. weissflogii* did exhibit significant changes to C:N ratios with Kyocera being highest and Fluorescent being lowest (Fig. 2L, Table S3) (One Way ANOVA, p < 0.05).

### Primary Productivity

Changes in chlorophyll normalized rates of PP were observed across light treatments and species and are presented in Figure 3 and in Table S4. In *P. marinus,* both oxygen and ^14^C derived estimates significantly differed. For ^14^C derived estimates, Fluorescent and Kyocera acclimated cells had PP rates 3% and 24% lower and Bright White acclimated cells had 35% lower PP rates compared to the Aquarium acclimated cells (Fig. 3A, Table S4) (One Way ANOVA p < 0.05). In O_2_ derived PP measurements, the general pattern remained the same, but cells acclimated to Fluorescent and Kyocera lights had 15% lower rates and Bright White acclimated cells having 35% lower PP rates compared to the Aquarium light (Fig. 3A, Table S4) (One Way ANOVA p < 0.05). In *Synechococcus sp.,* no significant differences between treatments in ^14^C PP measurements were measured (Fig. 3B, Table S4) (One Way ANOVA p > 0.05). In the O_2_ derived PP measurements cells acclimated to Aquarium had the highest rates, with Fluorescent, Kyocera and Bright White acclimated cells having rates 31, 58, and 44% lower compared to the Aquarium acclimation, respectively (Fig. 3B, Table S4) (One Way ANOVA p < 0.05). In *T. weissflogii,* PP rates did not significantly change among light treatments for either ^14^C or O_2_ based estimates (Fig. 3C, Table S4) (One Way ANOVA p > 0.05). Changes in rates for the O_2_ derived estimates were highest in the Aquarium treatment with the Kyocera, Bright White, and Fluorescent acclimated cells having 14, 19, and 17% lower rates respectively. A similar pattern was seen in the ^14^C derived estimates, with the Kyocera, Bright White, and Fluorescent acclimated cells having rates 17, 33, and 16% lower than the Aquarium acclimated cells (Fig. 3B, Table S4) (One Way ANOVA p > 0.05). Please note that rates of O_2_ evolution and C-fixation, while defined as net productivity, are biased by possible enhanced respiration rates during the light for the O_2_ experiments and also affected by the fact that ^14^C PP estimates descried rates between net and gross primary production.

**Figure 3:**
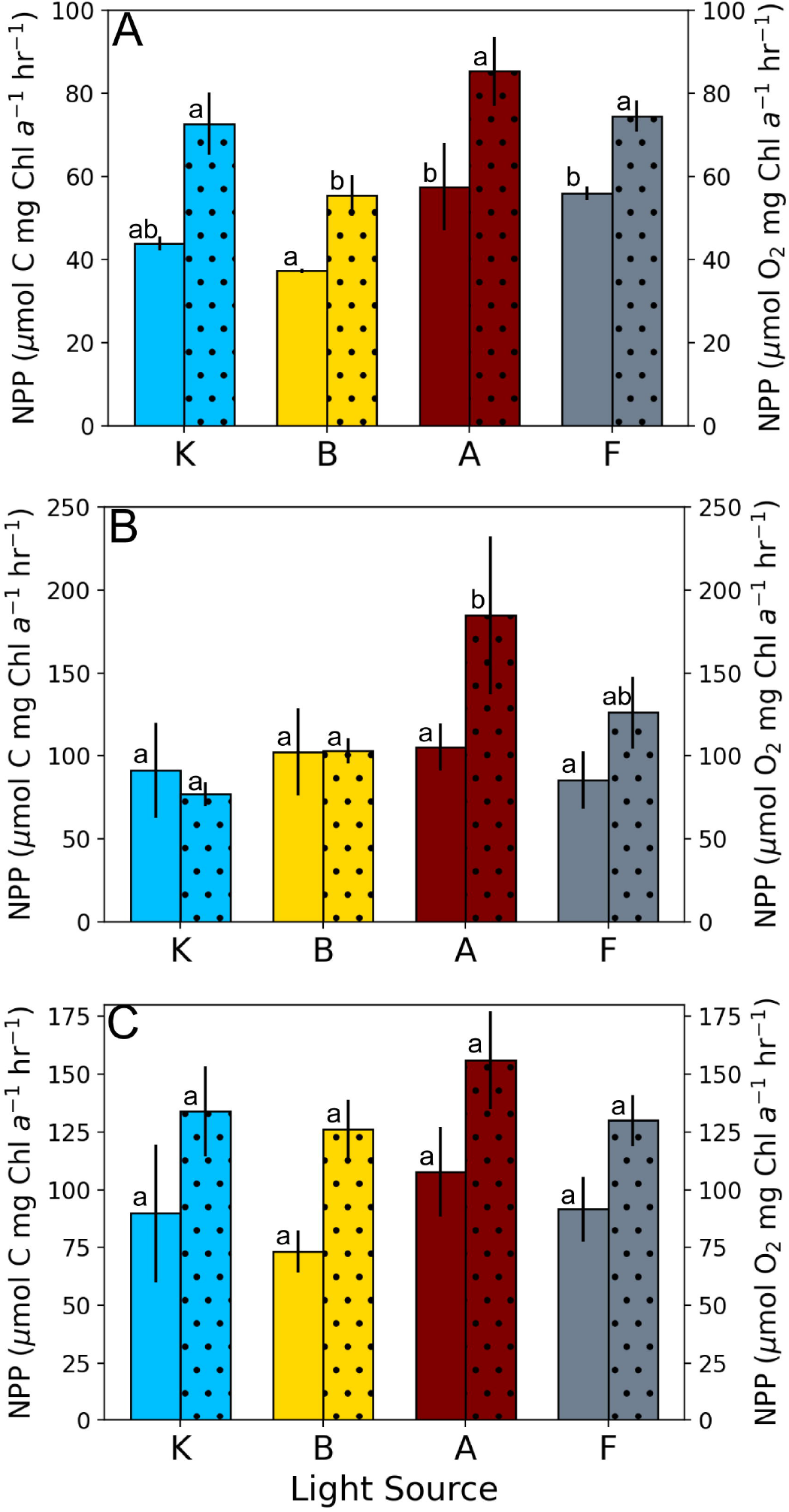
PP estimates from ^14^C and O_2_ incubations in units of *μ* mol C mg Chl *a*^-1^ hr^-1^ and *μ* mol O_2_ mg Chl *a*^-1^ hr^-1^, respectively for *Prochlorococcus marinus* (A), *Synechococcus sp.* (B), and *Thalassiosira weissflogii* (C). Data from ^14^C incubations are shown in solid bars while data from O_2_ incubations are in the dotted bars. Values shown are mean values for biological replicates ± s.d, n ≥ 2. Letters represent significant groupings from one-way ANOVA and HSD post hoc tests (p < 0.05). Note that the statistical groupings or the ^14^C and O_2_ incubations only refer to their respective measurements.

### Photophysiology

Photophysiology was measured to acquire additional parameters on photoacclimation and productivity (Fig. 4, Table S5). Please note that the spectrum and intensities of light in the FRRF differ from the spectrum of light in the acclimation as well as the light used to measure ^14^C and O_2_ PP. Photosynthesis vs. irradiance (P v I) curves are shown in Fig. 4 A, F, and K. Photophysiological data differed across species and light treatments. The α values for *P. marinus* were lower in the Kyocera and Aquarium treatment compared to the Bright White treatment while the Fluorescent treatment did not show significant differences to all other treatments (Fig. 4B, Table S5). (One Way ANOVA, p < 0.05). E_K_ and P_max_ values were significantly lower in the Bright White treatment than the Kyocera and Aquarium treatment, but no group was significantly different from the Fluorescent acclimation (Fig. 4C, D, Table S5) (One Way ANOVA, < 0.05). Non-photochemical quenching (NSV) did not significantly differ between treatments (Fig. 4E, Table S5) (One Way ANOVA, p > 0.05). No significant changes were seen in any FRRf derived photophysiological parameters for *Synechococcus sp.* (Fig. 4G, H, I, J, Table S5) (One Way ANOVA, p > 0.05). Similarly, no significant changes were seen in fit P v I curve statistics for *T. weissflogii* with the exception of the Fluorescent acclimated cells showing a significantly lower E_K_ value than all other acclimations (Fig. 4L. M, N, Table S5) (One Way ANOVA, p < 0.05). However, at acclimation light, the Fluorescent light acclimated cells showed significantly higher non-photochemical quenching (Fig. 4O, Table S5) (One Way ANOVA, p < 0.05).

**Figure 4:**
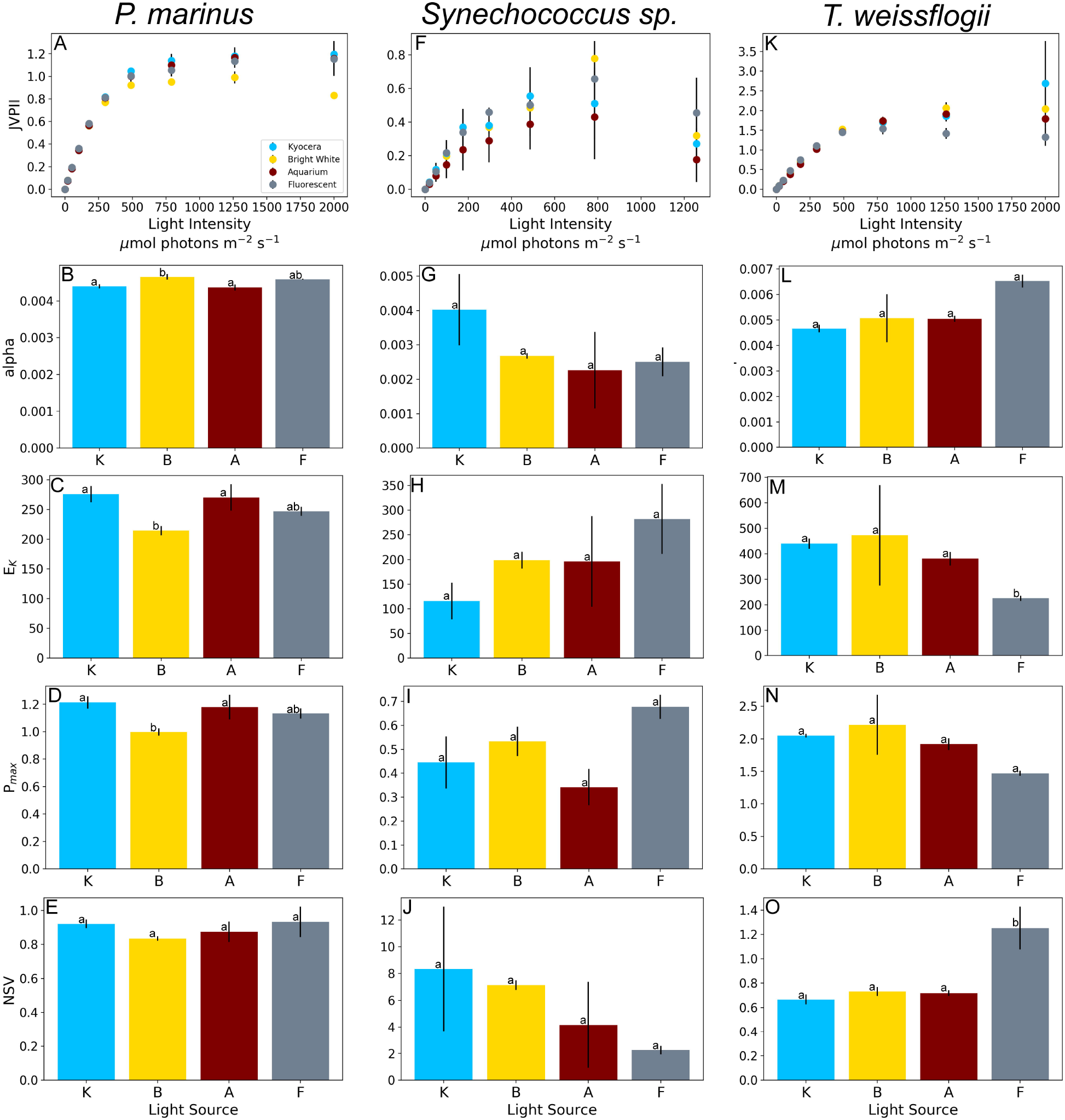
Photophysiological data derived from FRRf. JVPII (mol electron m^-3^ d^-1^) for *Prochlorococcus marinus* (A), *Synechococcus sp.* (F), and *Thalassiosira weissflogii* (K). α (mol electron μmol photons^-1^ m^-2^), E_K_ (μmol photons m^-2^ s^-1^), Pmax (mol electron m^-3^ d^-1^), as well as values for nonphotochemical quenching estimated as the normalized Stern-Volmer quenching coefficient (NSV) at acclimation light intensity for *Prochlorococcus marinus* (B-E), *Synechococcus sp.* (G-J), and *Thalassiosira weissflogii* (L-O). Values shown are mean values for biological replicates ± s.d, n ≥ 2. Letters represent significant groupings from one-way ANOVA and HSD post hoc tests (p < 0.05).

## Discussion

With the advent of cheap, readily accessible, and even customizable LED systems for scientific experiments, it is important to examine how the choice of growth light can influence ecologically and physiologically relevant cellular parameters. This is especially critical as we often attempt to predict how phytoplankton will respond to changes in environmental conditions (e.g. climate change) by using laboratory based experimental setups in which phytoplankton cultures are grown in different kind of temperature-controlled light-incubators. In this study we aimed to understand if different commercially available growth lights with differing light qualities but similar PAR affect phytoplankton growth, pigment content, elemental composition and productivity.

While light availability for photosynthesis is commonly regarded as photosynthetically active radiation (PAR) at wavelengths between 400 and 700 nm, phytoplankton cannot utilize each light wavelength equally (hence, PAR ≠ PUR). In our tested species, we determined that the Kyocera light supplied significantly higher usable photons over the absorbance range for the species *P. marinus* and *T. weissflogii,* while *Synechococcus* did not exhibit any PUR differences between the different lights tested (Fig. 1). Consequently, our expectation would be that both *P. marinus* and *T. weissflogii* would display larger differences while *Synechococcus* would show only minor responses. As presented in the results, we, however, found no changes in growth rate between the examined light sources for all cultures (Fig. 2, Table S3). This suggests that despite having different amounts of PUR, both *P. marinus* and *T. weissflogii* utilized a relatively similar amount of the emitted light energy to fuel growth while surplus energy consequently must either be utilized by other metabolic processes or dissipated. Our findings on growth concur with previous studies which have demonstrated that marine phytoplankton maintain constant growth rates under different spectral quality with seemingly similar PAR light intensities (Marchetti et al., 2013; Rivkin, 1989; Schellenberger Costa et al., 2013; Vadiveloo et al., 2015). While cells were maintaining their growth rates, the responses measured on cellular chl *a*, carbon and/or nitrogen content indicate that light sensing, harvesting and specific metabolic processes must have been affected by the changes in light spectrum.

Contrary to our expectations, some of the largest differences between light acclimations were seen in the *Synechococcus sp.* cultures. Despite having similar PUR, cellular chl *a* content, POC, and PON were significantly reduced under the Aquarium light (Fig. 2, Table S3). In addition, although not significantly different, a higher E_K_ and lower α was measured compared to the Kyocera light (Fig. 4, Table S5). We attribute some of the measured change to a lower absorbance of light in the in the blue 450-470 nm region and a higher absorbance in the yellow/green region where PE is strongly absorbing light (Fig. 1). Our spectral deconvolution identified higher PE/Chl *a* and PPC/Photoactive pigments ratios in the Aquarium acclimated cells compared to other treatments (Figure 1, Table S2). The dominance of PE in antenna complexes of phycobiliprotein containing cyanobacteria like *Synechococcus sp*. could potentially explain the discrepancy of a constant ^14^C PP while O_2_ PP was enhanced in the Kyocera compared to the Bright White LED acclimated cells. The different lighting conditions could cause PE to be associated with PSI rather than PSII, called state transition, leading to changes in electron flow through the photosystem. Under state II, PE would be associated to PSI and oxygen evolution will be reduced while electrons could be undergoing enhanced cyclic electron transport around PSI (Bailey et al., 2008). This would cause modification of the ratio of oxygen evolution to carbon fixation.

Based on most of the differences measured between the light conditions, it appears that *Synechococcus sp.* cells grown under the Aquarium light downregulate their photophysiology to a “high light” adapted state while maintaining growth and production. An additional change in either the state of photosystem II (PSII) or a change in the balance between PSII and PSI could have occurred explaining some of these responses. Photophysiological adjustments, not manifested in pigment content changes, could allow the cells to balance photosynthetically usable light absorption and alter linear and cyclic electron transport, which will ultimately change the cellular ratio of ATP to NADPH. Thus, the production of different metabolites could be regulated in these cells, leading to changes observed in cellular POC and PON. In all of our species tested, it is also possible that a small decrease in cell size could have caused changes of cellular chl *a* and POC/PON. For example, the strong decrease in chl *a* between Kyocera and Aquarium light in *Synechococcus sp.* could have been caused by a 25% decrease in cell diameter assuming a constant chl *a* cell^-1^ ratio. Spectral depended changes in cell size have been found for the green algae *Chlorella kessleri* (Koc et al. 2013) and *Chlorella vulgaris* (Lee and Palsson, 1996). While it is possible that cell size in our investigated species changed, we were unable to resolve these potentially small changes in cell diameter using the flow cytometer.

Concurrent to our expectations, *P. marinus* exhibited large differences in some of the cellular parameters when comparing Kyocera light (highest PUR) to the other lighting sources (Fig. 1, 2, Table S3). The enhanced PUR of the Kyocera light allowed higher production of cellular carbon and nitrogen in *P. marinus.* Interestingly, none of the further measurements (productivity and photophysiology) showed a strong response which could explain some of these data (Fig. 3,4 Table S4, S5). A reduced PPC/Photoactive pigment ratio was seen under the Bright White LED, which had the lowest PUR, however, a similar pigment ratio change was not seen in the Aquarium light which had similar PUR (Fig. 1, Table S2). Some of these differences and discrepancies presented could be caused by the differences of wavelength specific light utilization stemming from the different emissions, particularly in the 450 nm region where chl *a* absorbs, leading to altered metabolic base requirements (Fig. 1). Yet, as underlying mechanisms were not investigated in detail, no satisfying answer can be given at this moment.

Contrary to our expectations, no changes in PP (measured in either ^14^C or O_2_ incubations) were seen across light treatments in *T. weissflogii* (Fig. 3, Table S4). However, the changes in the C:N ratios of *T. weissflogii* between the Kyocera and Fluorescent light (Fig. 2, Table S3) indicate that the light quality influenced the cellular composition, a response previously observed for marine phytoplankton (Marchetti et al., 2013). As indicated by Geider and La Roche (2002), changes to cellular carbon and nitrogen quotas can be caused by changes of relative amounts of pigments, carbohydrates, lipids, proteins, etc. in a cell. While macromolecular composition (e.g. protein, lipid content) was not measured in this study, previous work has demonstrated that light quality can significantly affect the relative amounts of these cellular components (e.g. Gorai et al., 2014; Marchetti et al., 2013; Rivkin, 1989; Sánchez-Saavedra and Voltolina, 1994; Vadiveloo et al., 2015 and others). For *T. weissflogii,* an additional explanation could be that the higher emission around 450 nm where chl *a* absorbs in the Aquarium light compared to the Fluorescent light might have led to the observed reduction in cellular chl *a*.

The photoacclimation state in our tested species did not follow a general response pattern for the chosen growth lights, but our data demonstrate that the cells are sensing and responding to changes in the spectral quality in many ways. Indeed, it has been demonstrated that light spectra, in specific the perception of blue light, can affect photoacclimation in the diatom *P. tricornutum* (Schellenberger Costa et al., 2013) and red and far red light have found to stimulate carotenoid production in *Dunaliella salina* and *D. bardawil,* respectively (Sánchez-Saavedra et al., 1996; Xu et al., 2013). As such, it would be surprising to not find photophysiological adjustments in this study.

In general, all pigments have been found to be regulated under various light conditions (MacIntyre et al., 2002). *P. marinus* showed the lowest ratios of PPC/Photoactive pigments under the Bright White LED light and *Synechococcus sp.* showed the lowest ratios under the Aquarium LED (Fig. 1, Table S2). Based on the calculated E_K_, NSV, and the lack of photoinhibition in our P vs. I curves at acclimation light, our cells appear to be grown under non-light stressed conditions (sub-saturating light). Please note that FRRf derived parameters are not fully comparable to the steady state photophysiology the cells express during the light phase and only can be used for a direct comparison of the “relaxed state” of the photosystem between experiments. A change in the PPC/Photoactive pigment ratio could also be caused as a response to the specific emission intensity at the xanthophyll specific absorbance wavelength. Bright

White and Aquarium LEDs, which have the lowest emission in the 450-500 nm region where carotenoids (and also chl *a*) absorbance is dominant could thus have played a role in regulation of these pigments which would indicate a general readiness to cope with rapid changes to higher light intensities expected in turbulent or shallow mixed layers of stratified waters (Giovagnetti et al., 2014; Lewis et al., 1984; MacIntyre et al., 2002).

## Conclusions

The results from this study show that the light quality, not just quantity, provided to phytoplankton for growth can influence their physiology. While spectral sensitivity has been tested previously in several studies, an analysis of commercially available growth lights has not been conducted. The responses to the different growth lights used in this study were minimal in growth and productivity, however, responses in cellular composition, pigmentation, and estimated P vs. I curve parameters were all unique. In this study, distinct changes in POC, PON, C:N ratios and pigments were observed for each species. This is notable as it demonstrates the influence of growth light spectra on cell composition, which is important to keep in mind when performing laboratory experiments to constrain the biochemical composition of phytoplankton and their impact on biogeochemical cycles. We believe that cells growing under a light spectrum with low emission in the region where photoactive pigments absorb “sense” that they are growing in a lower light environment and may thus adapt their physiology accordingly. These findings could be important as several commercially sold LED incubators use Bright White LED systems, which might not be an ideal spectrum for all cultured species. While our measured responses seemed to be minor in many ways, experiments which investigate diurnal cycles under variable light intensities (sinus shape day night cycles, variable fluctuating light condition) could result in phytoplankton species being high-light stressed under relative low PAR intensities. Consequently, paying attention to spectral matching of emission and absorbance spectra can prove valuable in photobioreactors cultivation, particularly when used for the production of a desired cellular product (e.g. Abiusi et al., 2014; Baer et al., 2016; Chen et al., 2010; Walter et al., 2011). Numerical models often use photophysiological parameters as inputs to determine phytoplankton productivity and distribution, yet, data on E_K_, Pmax, and α are hard to interpret as not only light quantity but also specific light quality seemed to be a driving factor regulating these parameters. Overall, this study demonstrates that one must take into account the spectra of the growth lights as well as the pigmentation of the individual phytoplankton species, not just the light intensity. A simple “blue light, red light” combination of LEDs does not necessarily improve productivity, yet if specified to the absorbance to the cultured species it might. When designing laboratory experiments to mimic natural conditions one should also focus on the “true” spectrum in the water-column which can be obtained by more complex custom-made LED systems.

## Supplementary Data

Methods: Model parametrization.

Figure S1: Raw absorption data, modeled background, and fitted spectra for *P. marinus.*

Figure S2: Raw absorption data, modeled background, and fitted spectra for *Synechococcus sp.*

Figure S2: Raw absorption data, modeled background, and fitted spectra for *T. weissflogii.*

Table S1: Pigment gaussian peak (GP) parameters.

Table S2: Pigment Ratios.

Table S3: Growth rates, POC and PON cell^-1^, C:N. mol:mol ratios, and Chl *a* cell^-1^.

Table S4: Primary productivity (PP) estimates from ^14^C and O_2_ incubations normalized to Chl *a* content.

Table S5: Values for Photosynthesis vs Irradiance curve fits using Webb et al 1974.

## Acknowledgements

This research did not receive monetary funding from third party agencies in the public, commercial, or not-for-profit sections, yet we received the Kyocera light module from Kyocera (USA). Declarations of interest: none.

## References

Abiusi, F., Sampietro, G., Marturano, G., Biondi, N., Rodolfi, L., D’Ottavio, M., Tredici, M.R., 2014. Growth, photosynthetic efficiency, and biochemical composition of *Tetraselmis suecica* F&M-M33 grown with LEDs of different colors: Growth of *Tetraselmis* With Different LED Colors. Biotechnol. Bioeng. 111, 956–964. https://doi.org/10.1002/bit.25014

Baer, S., Heining, M., Schwerna, P., Buchholz, R., Hübner, H., 2016. Optimization of spectral light quality for growth and product formation in different microalgae using a continuous photobioreactor. Algal Res. 14, 109–115. https://doi.org/10.1016/j.algal.2016.01.011

Bailey, S., Melis, A., Mackey, K.R., Cardol, P., Finazzi, G., van Dijken, G., Berg, G.M., Arrigo, K., Shrager, J., Grossman, A., 2008. Alternative photosynthetic electron flow to oxygen in marine Synechococcus. Biochim. Biophys. Acta BBA-Bioenerg. 1777, 269–276.

Boatman, T.G., Lawson, T., Geider, R.J., 2017. A Key Marine Diazotroph in a Changing Ocean: The Interacting Effects of Temperature, CO2 and Light on the Growth of Trichodesmium erythraeum IMS101. PLOS ONE 12, e1168796. https://doi.org/10.1371/journal.pone.0168796

Boyd, P.W., Strzepek, R., Fu, F., Hutchins, D.A., 2010. Environmental control of open-ocean phytoplankton groups: Now and in the future. Limnol. Oceanogr. 55, 1353–1376. https://doi.org/10.4319/lo.2010.55.3.1353

Brand, L.E., Guillard, R.R.L., 1981. The effects of continuous light and light intensity on the reproduction rates of twenty-two species of marine phytoplankton. J. Exp. Mar. Biol. Ecol. 50, 119–132. https://doi.org/10.1016/0022-0981(81)90045-9

Chen, H.-B., Wu, J.-Y., Wang, C.-F., Fu, C.-C., Shieh, C.-J., Chen, C.-L, Wang, C.-Y., Liu, Y.-C., 2010. Modeling on chlorophyll a and phycocyanin production by Spirulina platensis under various light-emitting diodes. Biochem. Eng. J. 53, 52–56. https://doi.org/10.1016/j.bej.2010.09.004

Dickman, E.M., Vanni, M.J., Horgan, M.J., 2006. Interactive effects of light and nutrients on phytoplankton stoichiometry. Oecologia 149, 676–689. https://doi.org/10.1007/s00442-006-0473-5

Falkowski, P.G., 1984. Physiological responses of phytoplankton to natural light regimes. J. Plankton Res. 6, 295–307. https://doi.org/10.1093/plankt/6.2.295

Falkowski, P.G., Owens, T.G., 1978. Effects of light intensity on photosynthesis and dark respiration in six species of marine phytoplankton. Mar. Biol. 45, 289–295. https://doi.org/10.1007/BF00391815

Feng, Y., Warner, M.E., Zhang, Y., Sun, J., Fu, F.-X., Rose, J.M., Hutchins, D.A., 2008. Interactive effects of increased pCO _2_, temperature and irradiance on the marine coccolithophore *Emiliania huxleyi* (Prymnesiophyceae). Eur. J. Phycol. 43, 87–98. https://doi.org/10.1080/09670260701664674

Finkel, Z.V., Quigg, A., Raven, J.A., Reinfelder, J.R., Schofield, O.E., Falkowski, P.G., 2006. Irradiance and the elemental stoichiometry of marine phytoplankton. Limnol. Oceanogr. 51, 2690–2701. https://doi.org/10.4319/lo.2006.51.6.2690

Gao, K., Xu, J., Gao, G., Li, Y., Hutchins, D.A., Huang, B., Wang, L., Zheng, Y., Jin, P., Cai, X., Häder, D.-P., Li, W., Xu, K., Liu, N., Riebesell, U., 2012. Rising CO_2_ and increased light exposure synergistically reduce marine primary productivity. Nat. Clim. Change. https://doi.org/10.1038/nclimatel507

Geider, R., La Roche, J., 2002. Redfield revisited: variability of C:N:P in marine microalgae and its biochemical basis. Eur. J. Phycol. 37, 1–17. https://doi.org/10.1017/S0967026201003456

Giovagnetti, V., Flori, S., Tramontano, F., Lavaud, J., Brunet, C., 2014. The Velocity of Light Intensity Increase Modulates the Photoprotective Response in Coastal Diatoms. PLoS ONE 9, e103782. https://doi.org/10.1371/journal.pone.0103782

Glover, H.E., Keller, M.D., Spinrad, R.W., 1987. The effects of light quality and intensity on photosynthesis and growth of marine eukaryotic and prokaryotic phytoplankton clones. J. Exp. Mar. Biol. Ecol. 105, 137–159. https://doi.org/10.1016/0022-0981(87)90168-7

Gorai, T., Katayama, T., Obata, M., Murata, A., Taguchi, S., 2014. Low blue light enhances growth rate, light absorption, and photosynthetic characteristics of four marine phytoplankton species. J. Exp. Mar. Biol. Ecol. 459, 87–95. https://doi.org/10.1016/j.jembe.2014.05.013

Göritz, A., von Hoesslin, S., Hundhausen, F., Gege, P., 2017. Envilab: Measuring phytoplankton in-vivo absorption and scattering properties under tunable environmental conditions. Opt. Express 25, 25267. https://doi.org/10.1364/OE.25.025267

Healey, F.P., 1985. Interacting effects of light and nutrient limitation on the growth rate of Synechococcus linearis (Cyanophyceae)1. J. Phycol. 21, 134–146. https://doi.org/10.1111/j.0022-3646.1985.00134.x

Hessen, D.O., Leu, E., Færøvig, P.J., Falk Petersen, S., 2008. Light and spectral properties as determinants of C:N:P-ratios in phytoplankton. Deep Sea Res. Part II Top. Stud. Oceanogr. 55, 2169–2175. https://doi.org/10.1016/j.dsr2.2008.05.013

Hoppe, C.J.M., Holtz, L.-M., Trimborn, S., Rost, B., 2015. Ocean acidification decreases the light-use efficiency in an Antarctic diatom under dynamic but not constant light. New Phytol. 207, 159–171. https://doi.org/10.1111/nph.13334

Ivanikova, N.V., McKay, R.M.L., Bullerjahn, G.S., Sterner, R.W., 2007. Nitrate utilization by phytoplankton in Lake Superior is impaired by low nutrient (P, Fe) availability and seasonal light limitation - a cyanobacterial bioreporter study. J. Phycol. 43, 475–484. https://doi.org/10.1111/j.1529-8817.2007.00348.x

Kim, T.-H., Lee, Y., Han, S.-H., Hwang, S.-J., 2013. The effects of wavelength and wavelength mixing ratios on microalgae growth and nitrogen, phosphorus removal using Scenedesmus sp. for wastewater treatment. Bioresour. Technol. 130, 75–80. https://doi.org/10.1016/j.biortech.2012.11.134

Koc, C., Anderson, G.A., Kommareddy, A., 2013. Use of red and blue light-emitting diodes (LED) and fluorescent lamps to grow microalgae in a photobioreactor.

Kranz, S.A., Levitan, O., Richter, K.U., Prasil, O., Berman-Frank, I., Rost, B., 2010. Combined Effects of CO_2_ and Light on the N_2_-Fixing Cyanobacterium Trichodesmium IMS101: Physiological Responses. Plant Physiol. 154, 334–345.

Lee, C.-G., Palsson, B.Ø., 1996. Photoacclimation of Chlorella vulgaris to red light from light-emitting diodes leads to autospore release following each cellular division. Biotechnol. Prog. 12, 249–256.

Lehmuskero, A., Skogen Chauton, M., Boström, T., 2018. Light and photosynthetic microalgae: A review of cellular-and molecular-scale optical processes. Prog. Oceanogr. 168, 43–56. https://doi.org/10.1016/j.pocean.2018.09.002

Lewis, M., Horne, E., Cullen, J., Oakey, N., Platt, T., 1984. Turbulent motions may control phytoplankton photosynthesis in the upper ocean. Nature 311, 49–50.

MacIntyre, H.L., Kana, T.M., Anning, T., Geider, R.J., 2002. Photoacclimation of photosynthesis irradiance response curves and photosynthetic pigments in microalgae and cyanobacteria 1. J. Phycol. 38, 17–38.

Marchetti, J., Bougaran, G., Jauffrais, T., Lefebvre, S., Rouxel, C., Saint-Jean, B., Lukomska, E., Robert, R., Cadoret, J.P., 2013. Effects of blue light on the biochemical composition and photosynthetic activity of Isochrysis sp. (T-iso). J. Appl. Phycol. 25, 109–119. https://doi.org/10.1007/s10811-012-9844-y

Miao, H., Sun, L., Tian, Q., Wang, S., Wang, J., 2012. Study on the Effect of Monochromatic Light on the Growth of the Red Tide Diatom Skeletonema Costatum. Opt. Photonics J. 02, 152–156. https://doi.org/10.4236/opj.2012.23022

Morel, A., 1978. Available, usable, and stored radiant energy in relation to marine photosynthesis. Deep Sea Res. 25, 673–688. https://doi.org/10.1016/0146-6291(78)90623-9

Nielsen, E.S., 1952. The Use of Radio-active Carbon (C14) for Measuring Organic Production in the Sea. ICES J. Mar. Sci. 18, 117–140. https://doi.org/10.1093/icesjms/18.2.117

Oxborough, K., Moore, C.M., Suggett, D.J., Lawson, T., Chan, H.G., Geider, R.J., 2012. Direct estimation of functional PSII reaction center concentration and PSII electron flux on a volume basis: a new approach to the analysis of Fast Repetition Rate fluorometry (FRRf) data: Analysis of FRRf data: a new approach. Limnol. Oceanogr. Methods 10, 142–154. https://doi.org/10.4319/lom.2012.10.142

Prézelin, B., 1981. Light reactions in photosynthesis. Physiol. Bases Phytoplankton Ecol. 1–43.

Price, N.M., Harrison, G.I., Hering, J.G., Hudson, R.J., Nirel, P.M.V., Palenik, B., Morel, F.M.M., 1989. Preparation and Chemistry of the Artificial Algal Culture Medium Aquil. Biol. Oceanogr. 6, 443–461. https://doi.org/10.1080/01965581.1988.10749544

Redalje, D.G., Laws, E.A., 1983. The effects of environmental factors on growth and the chemical and biochemical composition of marine diatoms. I. Light and temperature effects. J. Exp. Mar. Biol. Ecol. 68, 59–79. https://doi.org/10.1016/0022-0981(83)90013-8

Rendon, S.M., Roldan, G.J.C., Voroney, R.P., 2013. Effect of carbon dioxide concentration on the growth response of Chlorella vulgaris under four different LED illumination. Int. J. Biotechnol. Wellness Ind. 2, 125–131.

Rhee, G.-ull, Gotham, I.J., 1981. The effect of environmental factors on phytoplankton growth: Light and the interactions of light with nitrate limitation1: Light-nutrient interactions. Limnol. Oceanogr. 26, 649–659. https://doi.org/10.4319/lo.1981.26.4.0649

Riebesell, U., Fabry, V.J., Hansson, L., Gattuso, J.-P., 2011. Guide to best practices for ocean acidification research and data reporting. Office for Official Publications of the European Communities.

Rivkin, R.B., 1989. Influence of irradiance and spectral quality on the carbon metabolism of phytoplankton. I. Photosynthesis, chemical composition and growth. Mar. Ecol. Prog. Ser. Oldendorf 55, 291–304.

Rost, B., Riebesell, U., Sültemeyer, D., 2006. Carbon acquisition of marine phytoplankton: effect of photoperiod length. Limnol. Oceanogr. 51, 12–20.

Sánchez-Saavedra, M. del P., Voltolina, D., 1994. The chemical composition of Chaetoceros sp. (Bacillariophyceae) under different light conditions. Comp. Biochem. Physiol. Part B Comp. Biochem. 107, 39–44. https://doi.org/10.1016/0305-0491(94)90222-4

Sánchez-Saavedra, M., Jiménez, C., Figueroa, F., 1996. Far-red light inhibits growth but promotes carotenoid accumulation in the green microalga Dunaliella bardawil. Physiol. Plant. 98, 419–423.

Schellenberger Costa, B., Jungandreas, A., Jakob, T., Weisheit, W., Mittag, M., Wilhelm, C., 2013. Blue light is essential for high light acclimation and photoprotection in the diatom Phaeodactylum tricornutum. J. Exp. Bot. 64, 483–493. https://doi.org/10.1093/jxb/ers340

Schulze, P.S.C., Barreira, L.A., Pereira, H.G.C., Perales, J.A., Varela, J.C.S., 2014. Light emitting diodes (LEDs) applied to microalgal production. Trends Biotechnol. 32, 422–430. https://doi.org/10.1016/j.tibtech.2014.06.001

Six, C., Finkel, Z.V., Irwin, A.J., Campbell, D.A., 2007. Light Variability Illuminates Niche-Partitioning among Marine Picocyanobacteria. PLoS ONE 2, e1341. https://doi.org/10.1371/journal.pone.0001341

Stadnichuk, I.N., Bulychev, A.A., Lukashev, E.P., Sinetova, M.P., Khristin, M.S., Johnson, M.P., Ruban, A.V., 2011. Far-red light-regulated efficient energy transfer from phycobilisomes to photosystem I in the red microalga Galdieria sulphuraria and photosystems-related heterogeneity of phycobilisome population. Biochim. Biophys. Acta BBA -Bioenerg. 1807, 227–235. https://doi.org/10.1016/j.bbabio.2010.10.018

Thrane, J.-E., Kyle, M., Striebel, M., Haande, S., Grung, M., Rohrlack, T., Andersen, T., 2015. Spectrophotometric Analysis of Pigments: A Critical Assessment of a High-Throughput Method for Analysis of Algal Pigment Mixtures by Spectral Deconvolution. PLOS ONE 10, eOl37645. https://doi.org/10.1371/journal.pone.0137645

Vadiveloo, A., Moheimani, N.R., Cosgrove, J.J., Bahri, P.A., Parlevliet, D., 2015. Effect of different light spectra on the growth and productivity of acclimated Nannochloropsis sp.(Eustigmatophyceae). Algal Res. 8, 121–127.

Walter, A., Carvalho, J.C. de, Soccol, V.T., Faria, A.B.B. de, Ghiggi, V., Soccol, C.R., 2011. Study of phycocyanin production from Spirulina platensis under different light spectra. Braz. Arch. Biol. Technol. 54, 675–682. https://doi.org/10.1590/S1516-89132011000400005

Wang, C.-Y., Fu, C.-C., Liu, Y.-C., 2007. Effects of using light-emitting diodes on the cultivation of Spirulina platensis. Biochem. Eng. J. 37, 21–25. https://doi.org/10.1016/j.bej.2007.03.004

White, E., Hoppe, C.J.M., Rost, B., 2020. The Arctic picoeukaryote Micromonas pusilla benefits from ocean acidification under constant and dynamic light. Biogeosciences 17, 635–647. https://doi.org/10.5194/bg-17-635-2020

Wilson, A., Punginelli, C., Gall, A., Bonetti, C., Alexandre, M., Routaboul, J.-M., Kerfeld, C.A., Van Grondelle, R., Robert, B., Kennis, J.T., others, 2008. A photoactive carotenoid protein acting as light intensity sensor. Proc. Natl. Acad. Sci. 105, 12075–12080.

Xu, B., Cheng, P., Yan, C., Pei, H., Hu, W., 2013. The effect of varying LED light sources and influent carbon/nitrogen ratios on treatment of synthetic sanitary sewage using Chlorella vulgaris. World J. Microbiol. Biotechnol. 29, 1289–1300. https://doi.org/10.1007/s11274-013-1292-6

