## Supplementary Information: Methods, Figures S1-S3, Tables S1-S5 for "Effects of spectral light quality on the growth, productivity, and elemental ratios in differently pigmented marine phytoplankton species"

List of Supplemental Information:

Methods: Model parametrization.

Figure S1: Raw absorption data, modeled background, and fitted spectra for *P. marinus.*

Figure S2: Raw absorption data, modeled background, and fitted spectra for *Synechococcus sp.*

Figure S2: Raw absorption data, modeled background, and fitted spectra for *T. weissflogii.*

Table S1: Pigment gaussian peak (GP) parameters.

Table S2: Pigment Ratios.

Table S3: Growth rates, POC and PON cell^-1^, C:N. mol:mol ratios, and Chl *a* cell^-1^.

Table S4: Primary productivity (PP) estimates from ^14^C and O_2_ incubations normalized to Chl *a* content.

Table S5: Values for Photosynthesis vs Irradiance curve fits using Webb et al 1974.

**Methods: Model Parameterization**

Gaussian peak (GP) parameters for individual pigments used in the spectral deconvolution model were fitted based on the reconstructed *in vivo* absorption scans from (Bidigare et al., 1990) for the following pigments: chlorophylls *a, b,* and *c*; photosynthetically active carotenoids, modeled by fucoxanthin (PSC); photoprotective carotenoids, modeled by beta-carotene (PPC); and phycoerythrin from *Synechococcus sp.* WH7803 (PE78). The *in vivo* absorption spectra provided by Bidigare et al. (1990) were further processed as required by Thrane et al., (2015) (set to zero absorption at 700 nm and normalized to their highest absorption peak). These normalized *in-vivo* absorption spectra were then processed by the R-script provided by Thrane et al., (2015) which recreates each pigment spectrum as a weighted sum of given GPs. Since species specific pigment-protein complexes induce shifts in pigment absorption characteristics, the model derived GP parameters were additionally modified to match the peak locations of our measured spectra. In addition, peak locations were cross referenced to peak maxima reported in literature (e.g. Murphy et al., 2017). Please note that pigment libraries used to determine the relative amount of pigments of each species were constrained to pigments known to be found in each species. Table S1 lists the pigment parameters used for each species.

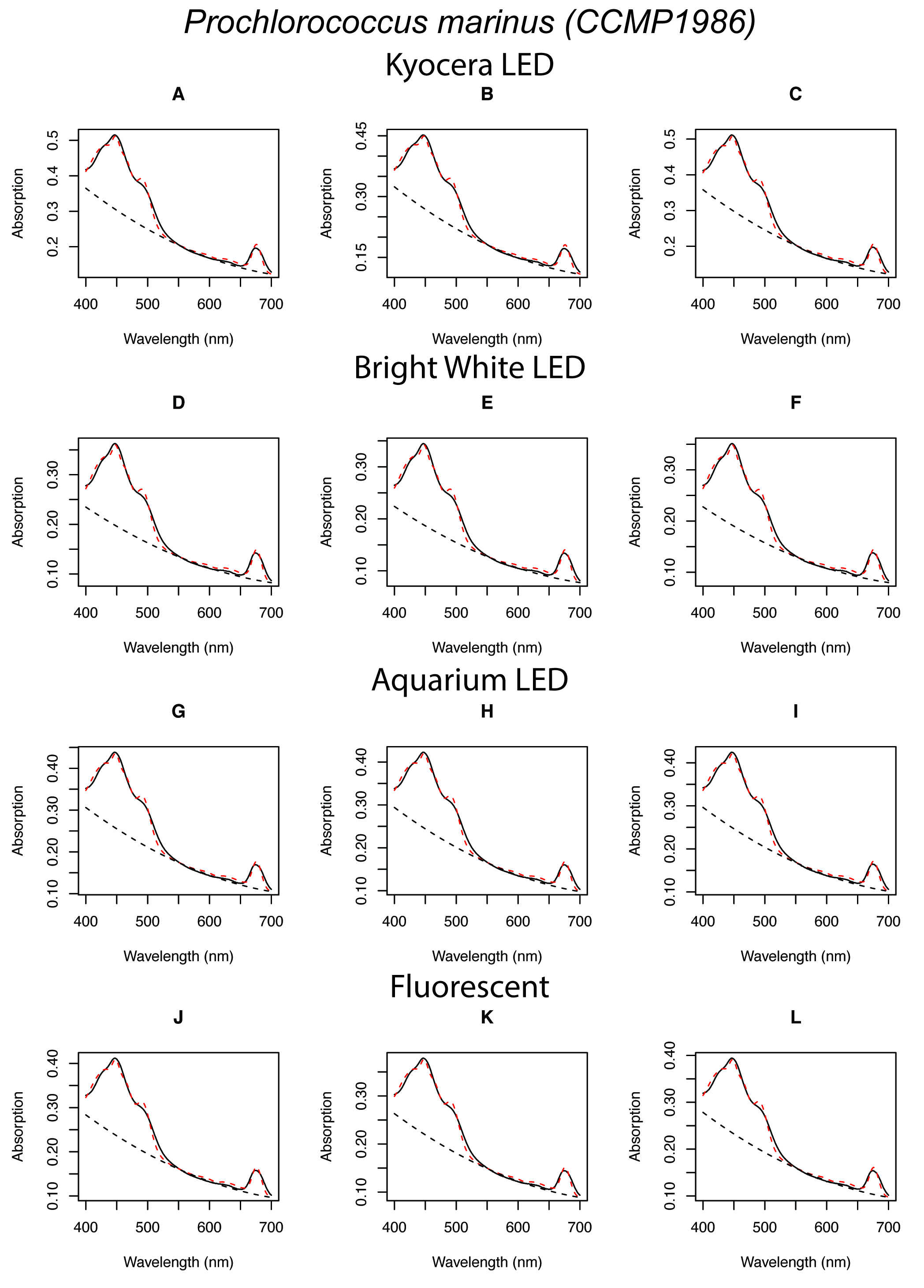

Figure S1: Raw absorption data, modeled background, and fitted spectra for *P. marinus.* Solid black lines show raw culture scan data, dashed black lines show modeled background, and dashed red lines show the fitted absorption spectra. Light acclimations are separated by row with Kyocera acclimations biological replicates being graphs A, B, C, and thus forth for the other 3 light treatments.

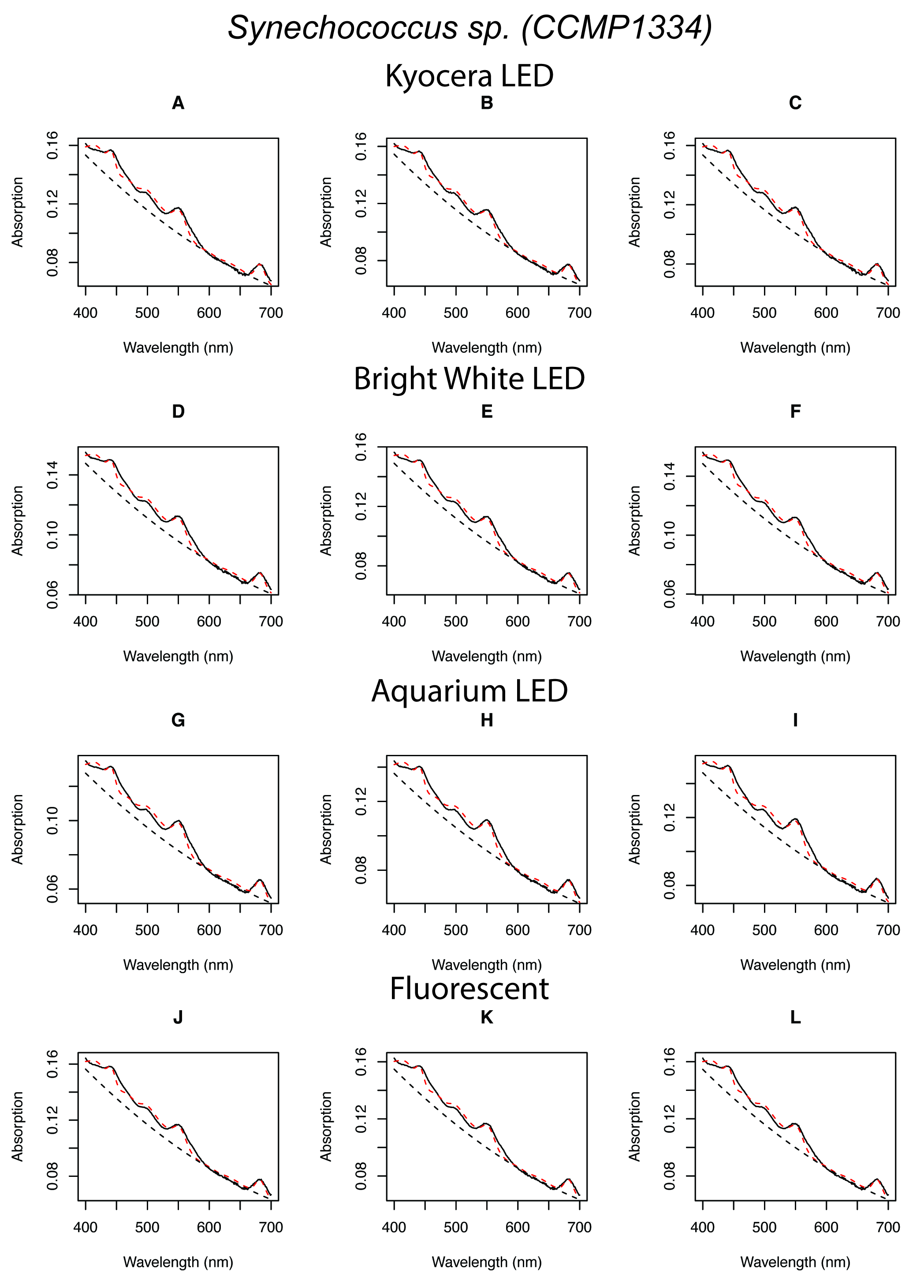

Figure S2: Raw absorption data, modeled background, and fitted spectra for *Synechococcus sp.* Solid black lines show raw culture scan data, dashed black lines show modeled background, and dashed red lines show the fitted absorption spectra. Light acclimations are separated by row with Kyocera acclimations biological replicates being graphs A, B, C, and thus forth for the other 3 light treatments.

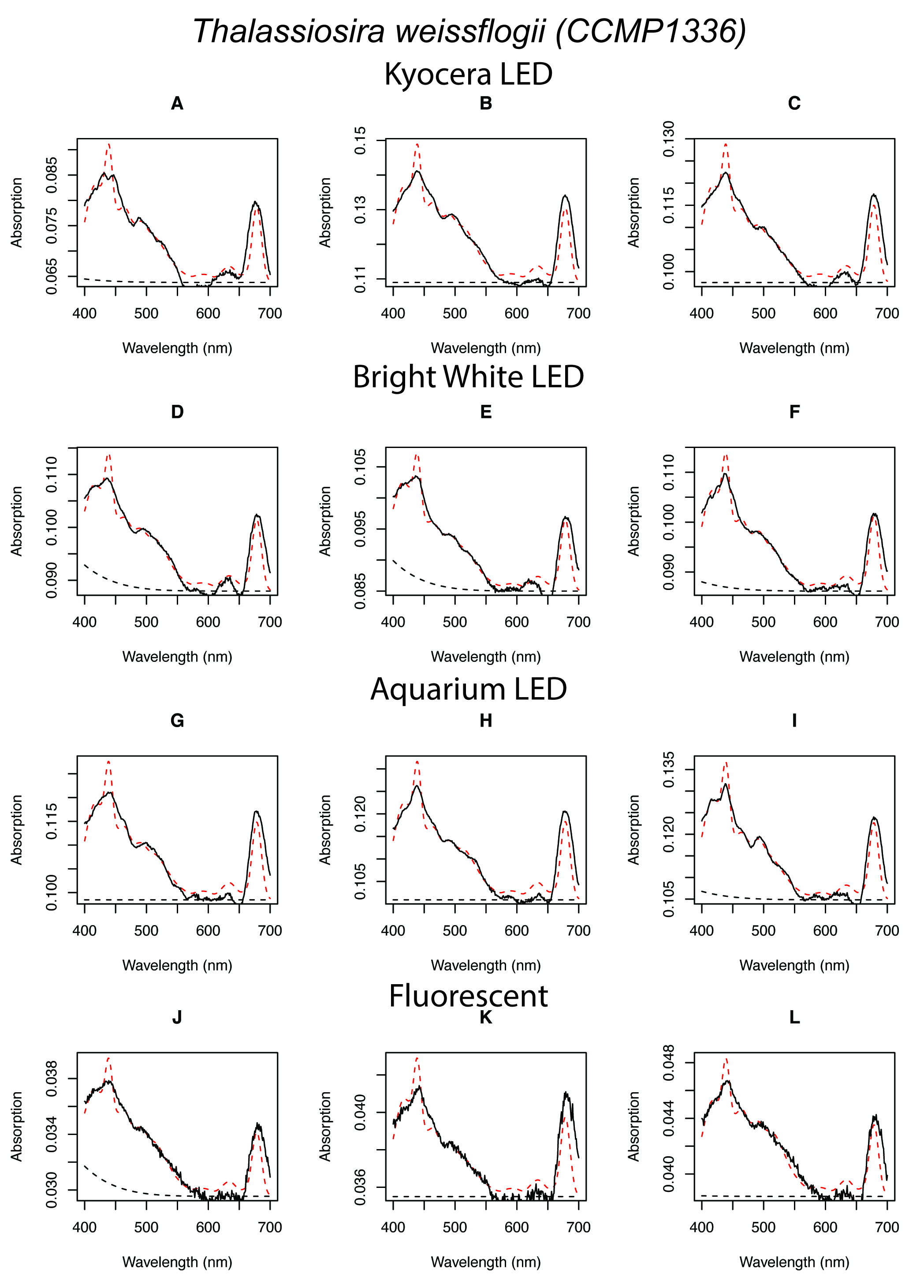

Figure S2: Raw absorption data, modeled background, and fitted spectra for *T. weissflogii*. Solid black lines show raw culture scan data, dashed black lines show modeled background, and dashed red lines show the fitted absorption spectra. Light acclimations are separated by row with Kyocera acclimations biological replicates being graphs A, B, C, and thus forth for the other 3 light treatments.

Table S1: Modified pigment gaussian peak (GP) parameters.

| Species | ***Prochlorochoccus marinus*** | | | ***Synechococcus sp.*** | | | ***Thalassiosira weissflogii*** | | |
| --- | --- | --- | --- | --- | --- | --- | --- | --- | --- |
| Pigment | Peak wave-length | Peak width | Peak height | Peak wave-length | Peak width | Peak height | Peak wave- length | Peak width | Peak height |
| Chl a | 427.5 | 16.8 | 0.71 | 421.5 | 16.8 | 0.71 | 418.5 | 16.8 | 0.71 |
|  | 450.0 | 6.6 | 0.70 | 444.0 | 6.6 | 0.70 | 441.0 | 6.6 | 0.70 |
|  | 460.3 | 5.0 | 0.00 | 454.3 | 5.0 | 0.00 | 451.3 | 5.0 | 0.00 |
|  | 479.0 | 4.6 | 0.00 | 473.0 | 4.6 | 0.00 | 470.0 | 4.6 | 0.00 |
|  | 496.5 | 4.6 | 0.00 | 490.5 | 4.6 | 0.00 | 487.5 | 4.6 | 0.00 |
|  | 514.0 | 4.6 | 0.00 | 508.0 | 4.6 | 0.00 | 505.0 | 4.6 | 0.00 |
|  | 531.5 | 4.6 | 0.00 | 525.5 | 4.6 | 0.00 | 522.5 | 4.6 | 0.00 |
|  | 549.2 | 3.7 | 0.00 | 543.2 | 3.7 | 0.00 | 540.2 | 3.7 | 0.00 |
|  | 557.0 | 6.0 | 0.00 | 563.0 | 6.0 | 0.00 | 559.0 | 6.0 | 0.00 |
|  | 583.7 | 8.2 | 0.02 | 589.7 | 8.2 | 0.02 | 585.7 | 8.2 | 0.02 |
|  | 598.1 | 9.8 | 0.05 | 604.1 | 9.8 | 0.05 | 600.1 | 9.8 | 0.05 |
|  | 624.9 | 9.6 | 0.08 | 630.9 | 9.6 | 0.08 | 626.9 | 9.6 | 0.08 |
|  | 641.0 | 10.8 | 0.09 | 647.0 | 10.8 | 0.09 | 643.0 | 10.8 | 0.09 |
|  | 658.2 | 5.0 | 0.04 | 664.2 | 5.0 | 0.04 | 660.2 | 5.0 | 0.04 |
|  | 669.7 | 12.8 | 0.00 | 675.7 | 12.8 | 0.00 | 671.7 | 12.8 | 0.00 |
|  | 678.0 | 8.5 | 0.75 | 684.0 | 8.5 | 0.75 | 680.0 | 8.5 | 0.75 |
|  | 701.3 | 4.7 | 0.00 | 707.3 | 4.7 | 0.00 | 703.3 | 4.7 | 0.00 |
|  | 719.0 | 4.6 | 0.00 | 725.0 | 4.6 | 0.00 | 721.0 | 4.6 | 0.00 |
|  | 736.5 | 4.6 | 0.00 | 742.5 | 4.6 | 0.00 | 738.5 | 4.6 | 0.00 |
| Chl b | 339.1 | 18.3 | 15.69 | N/A | N/A | N/A | N/A | N/A | N/A |
|  | 437.3 | 33.0 | 0.11 | N/A | N/A | N/A | N/A | N/A | N/A |
|  | 445.4 | 9.2 | 0.26 | N/A | N/A | N/A | N/A | N/A | N/A |
|  | 470.5 | 9.1 | 0.91 | N/A | N/A | N/A | N/A | N/A | N/A |
|  | 485.1 | 6.9 | 0.12 | N/A | N/A | N/A | N/A | N/A | N/A |
|  | 495.5 | 7.3 | 0.03 | N/A | N/A | N/A | N/A | N/A | N/A |
|  | 515.9 | 15.0 | 0.01 | N/A | N/A | N/A | N/A | N/A | N/A |
|  | 533.2 | 12.8 | 0.00 | N/A | N/A | N/A | N/A | N/A | N/A |
|  | 551.6 | 14.5 | 0.03 | N/A | N/A | N/A | N/A | N/A | N/A |
|  | 569.9 | 16.1 | 0.00 | N/A | N/A | N/A | N/A | N/A | N/A |
|  | 580.7 | 16.3 | 0.04 | N/A | N/A | N/A | N/A | N/A | N/A |
|  | 600.7 | 7.5 | 0.02 | N/A | N/A | N/A | N/A | N/A | N/A |
|  | 619.6 | 19.4 | 0.06 | N/A | N/A | N/A | N/A | N/A | N/A |
|  | 643.7 | 8.7 | 0.05 | N/A | N/A | N/A | N/A | N/A | N/A |
|  | 652.8 | 7.8 | 0.29 | N/A | N/A | N/A | N/A | N/A | N/A |
|  | 669.3 | 7.0 | 0.03 | N/A | N/A | N/A | N/A | N/A | N/A |
|  | 682.2 | 5.5 | 0.00 | N/A | N/A | N/A | N/A | N/A | N/A |
|  | 700.0 | 4.4 | 0.00 | N/A | N/A | N/A | N/A | N/A | N/A |
|  | 716.7 | 4.4 | 0.00 | N/A | N/A | N/A | N/A | N/A | N/A |
|  | 733.3 | 4.4 | 0.00 | N/A | N/A | N/A | N/A | N/A | N/A |
| Chl c | N/A | N/A | N/A | N/A | N/A | N/A | 405.4 | 24.7 | 0.29 |
|  | N/A | N/A | N/A | N/A | N/A | N/A | 445.4 | 13.3 | 0.40 |
|  | N/A | N/A | N/A | N/A | N/A | N/A | 454.5 | 6.9 | 0.26 |
|  | N/A | N/A | N/A | N/A | N/A | N/A | 467.2 | 9.8 | 0.74 |
|  | N/A | N/A | N/A | N/A | N/A | N/A | 484.0 | 11.2 | 0.03 |
|  | N/A | N/A | N/A | N/A | N/A | N/A | 495.8 | 13.3 | 0.02 |
|  | N/A | N/A | N/A | N/A | N/A | N/A | 516.1 | 8.9 | 0.02 |
|  | N/A | N/A | N/A | N/A | N/A | N/A | 533.3 | 7.8 | 0.02 |
|  | N/A | N/A | N/A | N/A | N/A | N/A | 551.3 | 8.9 | 0.03 |
|  | N/A | N/A | N/A | N/A | N/A | N/A | 566.9 | 6.9 | 0.03 |
|  | N/A | N/A | N/A | N/A | N/A | N/A | 585.0 | 9.5 | 0.08 |
|  | N/A | N/A | N/A | N/A | N/A | N/A | 599.0 | 8.9 | 0.02 |
|  | N/A | N/A | N/A | N/A | N/A | N/A | 620.2 | 11.3 | 0.05 |
|  | N/A | N/A | N/A | N/A | N/A | N/A | 636.5 | 7.0 | 0.12 |
|  | N/A | N/A | N/A | N/A | N/A | N/A | 653.0 | 7.5 | 0.02 |
|  | N/A | N/A | N/A | N/A | N/A | N/A | 663.3 | 8.4 | 0.01 |
|  | N/A | N/A | N/A | N/A | N/A | N/A | 682.6 | 4.4 | 0.00 |
|  | N/A | N/A | N/A | N/A | N/A | N/A | 700.0 | 4.4 | 0.00 |
|  | N/A | N/A | N/A | N/A | N/A | N/A | 716.7 | 4.4 | 0.00 |
|  | N/A | N/A | N/A | N/A | N/A | N/A | 733.3 | 4.4 | 0.00 |
| PSC | N/A | N/A | N/A | N/A | N/A | N/A | 406.2 | 33.6 | 0.14 |
|  | N/A | N/A | N/A | N/A | N/A | N/A | 432.5 | 14.0 | 0.16 |
|  | N/A | N/A | N/A | N/A | N/A | N/A | 444.6 | 9.3 | 0.12 |
|  | N/A | N/A | N/A | N/A | N/A | N/A | 463.1 | 12.6 | 0.48 |
|  | N/A | N/A | N/A | N/A | N/A | N/A | 486.7 | 14.8 | 0.52 |
|  | N/A | N/A | N/A | N/A | N/A | N/A | 498.8 | 15.2 | 0.50 |
|  | N/A | N/A | N/A | N/A | N/A | N/A | 516.7 | 10.5 | 0.38 |
|  | N/A | N/A | N/A | N/A | N/A | N/A | 531.4 | 12.5 | 0.43 |
|  | N/A | N/A | N/A | N/A | N/A | N/A | 550.9 | 12.6 | 0.16 |
|  | N/A | N/A | N/A | N/A | N/A | N/A | 570.0 | 14.3 | 0.05 |
|  | N/A | N/A | N/A | N/A | N/A | N/A | 587.1 | 11.9 | 0.01 |
|  | N/A | N/A | N/A | N/A | N/A | N/A | 604.9 | 7.7 | 0.01 |
|  | N/A | N/A | N/A | N/A | N/A | N/A | 619.3 | 4.6 | 0.00 |
|  | N/A | N/A | N/A | N/A | N/A | N/A | 636.3 | 4.4 | 0.00 |
|  | N/A | N/A | N/A | N/A | N/A | N/A | 653.0 | 4.4 | 0.00 |
|  | N/A | N/A | N/A | N/A | N/A | N/A | 669.7 | 4.4 | 0.00 |
|  | N/A | N/A | N/A | N/A | N/A | N/A | 686.3 | 4.4 | 0.00 |
|  | N/A | N/A | N/A | N/A | N/A | N/A | 703.0 | 4.4 | 0.00 |
|  | N/A | N/A | N/A | N/A | N/A | N/A | 719.7 | 4.4 | 0.00 |
|  | N/A | N/A | N/A | N/A | N/A | N/A | 736.3 | 4.4 | 0.00 |
| PE | N/A | N/A | N/A | 520.3 | 29.0 | 0.51 | N/A | N/A | N/A |
|  | N/A | N/A | N/A | 554.2 | 10.9 | 0.77 | N/A | N/A | N/A |
|  | N/A | N/A | N/A | 576.8 | 21.5 | 0.00 | N/A | N/A | N/A |
|  | N/A | N/A | N/A | 640.3 | 17.5 | 0.00 | N/A | N/A | N/A |
|  | N/A | N/A | N/A | 698.7 | 17.5 | 0.00 | N/A | N/A | N/A |
| PPC | 421.1 | 19.1 | 0.36 | 426.1 | 19.1 | 0.36 | 422.1 | 19.1 | 0.36 |
|  | 437.1 | 9.0 | 0.23 | 442.1 | 9.0 | 0.23 | 438.1 | 9.0 | 0.23 |
|  | 453.5 | 13.4 | 0.32 | 458.5 | 13.4 | 0.32 | 454.5 | 13.4 | 0.32 |
|  | 467.3 | 12.4 | 0.74 | 472.3 | 12.4 | 0.74 | 468.3 | 12.4 | 0.74 |
|  | 490.0 | 9.5 | 0.19 | 495.0 | 9.5 | 0.19 | 491.0 | 9.5 | 0.19 |
|  | 496.9 | 12.2 | 0.63 | 501.9 | 12.2 | 0.63 | 497.9 | 12.2 | 0.63 |
|  | 518.7 | 13.8 | 0.04 | 523.7 | 13.8 | 0.04 | 519.7 | 13.8 | 0.04 |
|  | 533.7 | 7.8 | 0.00 | 538.7 | 7.8 | 0.00 | 534.7 | 7.8 | 0.00 |
|  | 551.1 | 4.9 | 0.00 | 556.1 | 4.9 | 0.00 | 552.1 | 4.9 | 0.00 |
|  | 568.7 | 4.4 | 0.00 | 573.7 | 4.4 | 0.00 | 569.7 | 4.4 | 0.00 |
|  | 585.3 | 4.4 | 0.00 | 590.3 | 4.4 | 0.00 | 586.3 | 4.4 | 0.00 |
|  | 602.0 | 4.4 | 0.00 | 607.0 | 4.4 | 0.00 | 603.0 | 4.4 | 0.00 |
|  | 618.7 | 4.4 | 0.00 | 623.7 | 4.4 | 0.00 | 619.7 | 4.4 | 0.00 |
|  | 635.3 | 4.4 | 0.00 | 640.3 | 4.4 | 0.00 | 636.3 | 4.4 | 0.00 |
|  | 652.0 | 4.4 | 0.00 | 657.0 | 4.4 | 0.00 | 653.0 | 4.4 | 0.00 |
|  | 668.7 | 4.4 | 0.00 | 673.7 | 4.4 | 0.00 | 669.7 | 4.4 | 0.00 |
|  | 685.3 | 4.4 | 0.00 | 690.3 | 4.4 | 0.00 | 686.3 | 4.4 | 0.00 |
|  | 702.0 | 4.4 | 0.00 | 707.0 | 4.4 | 0.00 | 703.0 | 4.4 | 0.00 |
|  | 718.7 | 4.4 | 0.00 | 723.7 | 4.4 | 0.00 | 719.7 | 4.4 | 0.00 |
|  | 735.3 | 4.4 | 0.00 | 740.3 | 4.4 | 0.00 | 736.3 | 4.4 | 0.00 |

Table S2: Pigment Ratios for *P. marinus, Synechococcus sp.,* and *T. weissflogii*. Values shown are mean values for biological replicates ± s.d, n = 3. Letters represent significant groupings from one-way ANOVA and HSD post hoc tests (p < 0.05).

|  | | Kyocera | Bright White | Aquarium | Fluorescent |
| --- | --- | --- | --- | --- | --- |
| *Prochlorococcus marinus* | Chl *b* / Chl *a* | 0.069 ± 0.004 ^a^ | 0.070 ± 0.002 ^a^ | 0.067 ± 0.001 ^a^ | 0.064 ± 0.001 ^a^ |
|  | PPC / Chl *a* | 0.503 ± 0.003 ^a^ | 0.474 ± 0.003 **^b^** | 0.506 ± 0.003 ^a^ | 0.510 ± 0.002 ^a^ |
|  | PPC/Photoactive Pigments | 0.471 ± 0.001 ^a^ | 0.444 ± 0.002 ^b^ | 0.474 ± 0.003 ^a^ | 0.479 ± 0.002 ^a^ |
|  | PPC / Total Pigments | 0.320 ± 0.001 ^a^ | 0.307 ± 0.001 ^b^ | 0.322 ± 0.002 ^a^ | 0.324 ± 0.001 ^a^ |
| *Synechococcus sp.* | PE / Chl *a* | 3.479 ± 0.048 ^a^ | 3.539 ± 0.054 ^a^ | 3.751 ± 0.021 ^b^ | 3.502 ± 0.019 ^a^ |
|  | PPC / Chl *a* | 0.194 ± 0.008 ^a^ | 0.191 ± 0.002 ^a^ | 0.158 ± 0.011 ^b^ | 0.204 ± 0.003 ^a^ |
|  | PPC/Photoactive Pigments | 0.043 ± 0.002 ^a^ | 0.042 ± 0.0003 ^a^ | 0.033 ± 0.002 ^b^ | 0.045 ± 0.001 ^a^ |
|  | PPC / Total Pigments | 0.042 ± 0.002 ^a^ | 0.040 ± 0.0003 ^a^ | 0.032 ± 0.002 ^b^ | 0.043 ± 0.001 ^a^ |
| *Thalassiosira weissflogii* | Chl *c* / Chl *a* | 0.077 ± 0.007 ^a^ | 0.064 ± 0.007 ^a^ | 0.068 ± 0.023 ^a^ | 0.086 ± 0.018 ^a^ |
|  | PSC / Chl *a* | 0.274 ± 0.047 ^a^ | 0.270 ± 0.047 ^a^ | 0.257 ± 0.035 ^a^ | 0.323 ± 0.103 ^a^ |
|  | PPC / Chl *a* | 0.033 ± 0.013 ^a^ | 0.028 ± 0.026 ^a^ | 0.026 ± 0.031 ^a^ | 0.022 ± 0.020 ^a^ |
|  | PPC/Photoactive Pigments | 0.024 ± 0.010 ^a^ | 0.021 ± 0.020 ^a^ | 0.021 ± 0.025 ^a^ | 0.017 ± 0.015 ^a^ |
|  | PPC / Total Pigments | 0.024 ± 0.010 ^a^ | 0.020 ± 0.019 ^a^ | 0.020 ± 0.023 ^a^ | 0.016 ± 0.015 ^a^ |

Table S3: Growth rates (*µ* d^-1^), POC and PON (fg cell^-1^ for *P. marinus and Synechococcus,* pg cell^-1^ for *T. weissflogii*) , C:N (mol:mol), and Chl *a* cell^-1^ (fg Chl *a* cell^-1^ for *P. marinus and Synechococcus,* pg Chl *a* cell^-1^ for *T. weissflogii*). Values shown are mean values for biological replicates ± s.d, n ≥ 2. Stars represent samples which have n=2 instead of n=3. Letters represent significant groupings from one-way ANOVA and HSD post hoc tests (p < 0.05).

|  | | Kyocera | Bright White | Aquarium | Fluorescent |
| --- | --- | --- | --- | --- | --- |
| *Prochlorococcus marinus* | Growth rate | 0.25 ± 0.04^a^ | 0.24 ± 0.04^a^ | 0.25 ± 0^a^ | 0.24 ± 0.04^a^ |
|  | POC | 99.2 ± 6.2^a^ | 55.5 ± 0.8^b^ | 73.9 ± 4.1^c^ | 79.8 ± 6.9^c^ |
|  | PON | 21.0 ± 1.1^a^ | 12.0 ± 0.4^b^ | 16.0 ± 1.0^c^ | 16.7 ± 0.9^c^ |
|  | C:N | 5.52 ± 0.38^a^ | 5.38 ± 0.09^a^ | 5.38 ± 0.05^a^ | 5.56 ± 0.19^a^ |
|  | Chl *a* | 2.88 ± 0.32^a^ | 2.43 ± 0.07^a^ | 2.63 ± 0.15^a^ | 2.76 ± 0.08^a^ |
| *Synechococcus sp.* | Growth rate | 0.35 ± 0.01^a^ | 0.33 ± 0.01^ab^ | 0.30 ± 0.01^b^ | 0.33± 0.01^ab^ |
|  | POC | 308.1 ± 10.0^a^ | 289.7 ± 13.0^a^ | 258.6 ± 3.7*^b^ | 304.9 ± 7.0^a^ |
|  | PON | 60.5 ± 1.9^a^ | 57.3 ± 2.5^a^ | 49.8 ± 1.6*^b^ | 57.9 ± 1.7^a^ |
|  | C:N | 5.95 ± 0.21^a^ | 5.89 ± 0.18^a^ | 6.07 ± 0.28*^a^ | 6.15 ± 0.15^a^ |
|  | Chl *a* | 5.06 ± 0.28^ab^ | 3.61 ± 0.71^ab^ | 2.82 ± 0.62^a^ | 5.20 ± 0.67^b^ |
| *Thalassiosira weissflogii* | Growth rate | 0.86 ± 0.05^a^ | 0.83 ± 0.04^a^ | 0.86 ± 0.03^a^ | 0.84 ± 0.03^a^ |
|  | POC | 93.7 ± 7.0^a^ | 86.2 ± 11.3^ab^ | 67.7 ± 2.2^b^ | 81.8 ± 11.7^ab^ |
|  | PON | 18.2 ± 2.3^a^ | 19.3 ± 1.2^a^ | 15.1 ± 1.0^a^ | 19.5 ± 2.4^a^ |
|  | C:N | 6.04 ± 0.32^a^ | 5.21 ± 0.39^ab^ | 5.26 ± 0.43^ab^ | 4.89 ± 0.22^b^ |
|  | Chl *a* | 6.85 ± 0.28^ab^ | 6.12 ± 0.49^ab^ | 5.19 ± 0.75^a^ | 7.11 ± 0.57^b^ |

Table S4: Primary productivity (PP) from ^14^C and O_2_ incubations normalized to Chl *a* content. Values shown are mean values for biological replicates ± s.d, n ≥ 2. Stars represent samples which have n=2 instead of n=3. Letters represent significant groupings from one-way ANOVA and HSD post hoc tests (p < 0.05) between the different light condition of each species.

|  | | Kyocera | Bright White | Aquarium | Fluorescent |
| --- | --- | --- | --- | --- | --- |
| *Prochlorococcus marinus* | ^14^C-PP  (*µ*mol C mg Chl *a*^-1^ hr^-1^) | 43.86 ± 1.66^ab^ | 37.39 ± 0.54^a^ | 57.57 ± 10.56*^b^ | 55.98 ± 1.71*^b^ |
|  | O_2_-PP  (*µ*mol O_2_ mg Chl *a*^-1^ hr^-1^) | 72.67 ± 7.52^a^ | 55.53 ± 4.88^b^ | 85.31 ± 8.41^a^ | 74.53 ± 3.71^a^ |
| *Synechococcus sp.* | ^14^C-PP  (*µ*mol C mg Chl *a*^-1^ hr^-1^) | 91.31 ± 28.38*^a^ | 102.39 ± 26.19^a^ | 105.20 ± 14.04^a^ | 85.48 ± 17.17^a^ |
|  | O_2_-PP  (*µ*mol O_2_ mg Chl *a*^-1^ hr^-1^) | 76.95 ± 7.24^a^ | 103.02 ± 7.74^a^ | 184.81 ± 47.36^b^ | 126.10 ± 21.90^ab^ |
| *Thalassiosira weissflogii* | ^14^C-PP  (*µ*mol C mg Chl *a*^-1^ hr^-1^) | 89.83 ± 29.83^a^ | 73.26 ± 8.96^a^ | 107.75 ± 19.41^a^ | 91.63 ± 14.00^a^ |
|  | O_2_-PP  (*µ*mol O_2_ mg Chl *a*^-1^ hr^-1^) | 133.89 ± 19.49^a^ | 126.05 ± 12.67^a^ | 156.14 ± 21.09^a^ | 130.00 ± 11.12^a^ |

Table S5: Photosynthesis vs Irradiance curve from FRRF measurements using the Webb et al 1974 fit parameters. Values shown are average values for biological replicates ± s.d, n ≥ 2. Stars represent samples which have n=2 instead of n=3. Letters represent significant groupings from one-way ANOVA and HSD post hoc tests (p < 0.05).

|  | | Kyocera | Bright White | Aquarium | Fluorescent |
| --- | --- | --- | --- | --- | --- |
| *Prochlorococcus marinus* | α (mol electron µmol photons^-1^ m^-2^) | 0.00440 ± 0.00006^a^ | 0.00465 ± 0.00008^b^ | 0.00436 ± 0.00008^a^ | 0.00459± 0.00001^ab^ |
|  | E_K_ (*µ*mol photons m^-2^ s^-1^) | 276 ± 14^a^ | 215 ± 8^b^ | 270 ± 22^a^ | 247 ± 8^ab^ |
|  | P_max_ (mol electrons m^-2^ d^-1^) | 1.213 ± 0.044^a^ | 0.998 ± 0.026^b^ | 1.179 ± 0.090^a^ | 1.134 ± 0.037^ab^ |
|  | NSV (at 100 µE) | 0.921 ± 0.025^a^ | 0.834 ± 0.013^a^ | 0.875 ± 0.060^a^ | 0.932 ± 0.090^a^ |
| *Synechococcus sp.* | α (mol electron µmol photons^-1^ m^-2^) | 0.00402 ± 0.00104^a^ | 0.00268 ± 0.00008*^a^ | 0.00226 ± 0.00111^a^ | 0.00251± 0.00041^a^ |
|  | E_K_ (*µ*mol photons m^-2^ s^-1^) | 116 ± 37^a^ | 198 ± 17*^a^ | 196 ± 92^a^ | 282 ± 7^b^ |
|  | P_max_ (mol electrons m^-2^ d^-1^) | 0.445 ± 0.110^a^ | 0.533 ± 0.061*^a^ | 0.342 ± 0.076^a^ | 0.678 ± 0.050^a^ |
|  | NSV (50 at µE) | 8.349 ± 4.658^a^ | 7.141 ± 0.358*^a^ | 4.151 ± 3.225^a^ | 2.244 ± 0.317^a^ |
| *Thalassiosira weissflogii* | α (mol electron µmol photons^-1^ m^-2^) | 0.00466 ± 0.00015*^a^ | 0.00507 ± 0.00094^a^ | 0.00504 ± 0.00012^a^ | 0.00653± 0.00026^a^ |
|  | E_K_ (*µ*mol photons m^-2^ s^-1^) | 440 ± 20*^a^ | 472 ± 197^a^ | 381 ± 27^a^ | 225 ± 10^b^ |
|  | P_max_ (mol electrons m^-2^ d^-1^) | 2.050 ± 0.029*^a^ | 2.214 ± 0.462^a^ | 1.917 ± 0.091^a^ | 1.468 ± 0.039^a^ |
|  | NSV (at 100 µE) | 0.666 ± 0.041^a^ | 0.733 ± 0.037^a^ | 0.718 ± 0.023^a^ | 1.252 ± 0.176^b^ |
